# Soil Microbial and Biochemical Properties under Conservation Agriculture in rice-based cropping systems in lower Indo-Gangetic Plain of West Bengal

**DOI:** 10.64898/2026.08.31.748290

**Authors:** Puja Singh, M Jaison, Niharendu Saha, Susanta Dutta, Aradhana Sen, Tufleuddin Biswas, Biswapati Mandal, Siddhartha Mukherjee, Bishnuprasad Dash, Biswabara Sahu, Ruby Patel, Amrita Dasgupta

## Abstract

Microbial and biochemical properties of soil respond quickly with change in management practices, compared to chemical and physical properties. Moreover, impact of conservation agriculture (CA) on soil microbial properties is limited to microbial enumeration, but its effect on soil enzyme and microbial activity is little documented. To address these problems soil enzyme activities [dehydrogenase (DHA), β-glucosidase (BGA), acid phosphatase (AcP) and alkaline phosphatase (AlP) and fluoresceine diacetate (FDA)], activities of soil microbes ((Nitrogen fixation (NFBAct), Phosphate solubilization (PSBAct) & Cellulolytic activities (CDBAct)), microbial biomass ((Soil microbial biomass carbon (SMBC) & soil microbial biomass nitrogen (SMBN)) and available nutrient were studied to evaluate biological soil health in alluvial soil of lower Indo-Gangetic plain (IGP) under different degree of CA. Field experiment was conducted in split plot design (SPD), under 3 cropping systems (RMCp: rice-maize-cowpea; RWGg: rice-wheat-green gram; RCfBr; rice-cauliflower-bororice/summer rice). Tillage operations (CT: conventional tillage; MT: minimum tillage and ZT: zero tillage) was main plot and residue application as sub plot treatments [(R_0_ (no residue), R_50_ (50% residue) and R_100_ (100% residue)]. All the treatments were replicated thrice. Considering values of biological soil health index (BSHI), it is found that among different degree of CA, ZT (0.464) and (MT=0.441) and R_100_ (0.464) treatment showed better response. Among different cropping system RMCp (0.359) & RWGg (0.343) outperformed RCfBr (0.609) cropping system with respect to (wrt) microbial and biochemical properties of the soil. Result indicated that for restoring microbial and biochemical properties of soil practices of CA can be promising solution.

**Importance:** ntensive farming is profit oriented and their sustainability are at risk. CA is low input farming and its practices (low mechanical disturbance, crop rotation, residues retention) help to restore agro-ecosystem. Thus, CA is emerging as potential technology to negate adverse impact of modern faming on soil. Soil microbial properties (population, activities, biomass and enzymes) are key indicators of soil health and quality as they are highly sensitive to management practices. Hence to address impact of CA practices on soil, microbial and biochemical aspects are studied. Degree of CA practice with higher BSHI can be popularized in filed to restore soil.

**Highlights:**

1. Actual response and activities of soil microbes and enzymes under CA were lacking.
2. Ranking of different treatments based on multivariate analysis and BSHI score indicated ZT + R_100_ was best suited to soil microbes and enzyme, while, higher fertilizer dose showed negative impact on them.
3. Higher SMB was detected in soil under conservation practices especially under RMCp & RWGg.
4. RCfBr showed poor performance as legume haven’t incorporated in rotation.
5. CT favour enzyme activities, especially when combined with residue application under legume-based cropping systems.

## 1. Introduction

Intensive agriculture practices are becoming threat to microbial and biochemical properties of the soil, which can be seen in term of decline in soil fertility and productivity. It is high time for agriculturists to implement soil management practices that sustain crop productivity along with sustaining food security and environmental (Singh et al. 2021). To address such issues related to soil properties CA is a potential technology. Principles of CA (a) Avoiding soil disturbance (to prevents short-term peak of biological activity associated with flushes of carbon (C) and nitrogen (N) loss; (b) Covering soil surface with residue (provide substrate for soil microbes); (c) crop diversification (with inclusion of legumes) can restore activities of soil microbes and enzymes (Leal et al. 2020; Pires et al. 2020). Such practices intensify C sequestration and stratification, improve soil structure and stability, augment substrate diversity, stabilize ecosystem, and above all reduce demographic stochasticity to restore microbial habitat (Passinato et al. 2021). Besides, soil productivity & fertility primarily depends on biological health, which includes soil microbes, their activities & biomass, enzymatic etc. These properties are highly dynamic and sensitive to the changes in soil attributes associated with soil management practices (Singh et al. 2024).

Habig et al. (2015) reported CA yield higher activities of soil microbes (e.g., FLNF, PSB and CDB). Parihar et al. (2016) observed a significant positive effect of CA practices on soil MBC (45-48.9% increase in 0–30 cm depth of a sandy loam soil) under maize-based rotation after 6 years of CA adoption. Enzymes activity is an excellent short-term indicator of soil biological and biochemical fertility as it reflects microbiological processes that mineralize organic nutrients for growth and development of plants (Passinato et al. 2021). Soil enzyme are “sensors” that integrate information from microbial status and physico-chemical condition of the soil (Wittmann et al. 2024). Among various soil enzymes dehydrogenase (DHA), β-glucosidase (BGA), acid (AcP) & alkaline phosphatase (AlP) etc. are considered important as involved in mineralization of C, N, and P. DHA is associated with viable cells and reflects oxidative activity of soil microbial population (Borase et al. 2020). AcP and AlP mediates release of inorganic phosphorus from organically-bound phosphorus added as litter or other organic debris (Zheng et al. 2021). FDA reflect strong correlation with soil microbial properties compared to DHA. Mendes et al. (2021) gave similar weights to BGA and DHA in soil nutrient cycling.

Intricacy of soil microbes and their function for sustenance of crop production under CA are poorly documented, due to poor understanding of soil microbial and biochemical properties under the microclimate created under CA. To our knowledge based on literature available, effect of CA practices on soil microbial world and associated activities and processes is scanty. Most of the research conducted on soil microbes under CA is confined to microbial enumeration and diversity study but activities of soil microbes and enzymes under altered soil conditions, are still rudimentary. The experiment was carried to study **‘Soil Microbial and Biochemical Properties under Conservation Agriculture in rice-based cropping systems in Lower Indo-Gangetic Plain of West Bengal’** with the hypothesis that microclimate created under CA favors the microbial and biochemical properties of soil. Individual soil parameters alone may not be sufficient for decision making regarding sustainability of the cropping system (Saha & Mandal, 2009). In CA based management systems in IGP, studies on various soil parameters especially physico-chemical properties and few reports on biological properties have been documented but in isolation. The main objective of this study was to assess soil microbial properties, through the activity of soil microbes and enzymes as bioindicators, along with their impact on each other, under CA in alluvial soil of lower IGP.

## 2. Materials and methods

### 2.1 Experimental Site

A field experiment was done during the Kharif (rainy) season of 2018 at Balindi Farm, as part of the Centre for Advanced Agriculture Science and Technology (CAAST) on Conservation Agriculture at Bidhan Chandra Krishi Vishwavidyalaya (22° 57’ 46” N, 88° 31’ 48”’ E, 9.75 m above mean sea level). The site is located in the New Alluvial Zone (NAZ) of the lower IGP, characterized by a subtropical humid climate with an average annual rainfall of 1500 mm. Minimum and maximum temperature averaged 12.5°C and 36.3°C, respectively, with a mean annual temperature of 32°C. Soil of this region belong to Inceptisol is having silty clay loam texture. Initial soil properties (depth 0–20cm) studied evidenced in Table 1.

**Table 1:** Initial soil properties of Balindi Farm of BCKV, Mohanpur:

| SI. No | Soil | Result | Method followed (Reference) |
| --- | --- | --- | --- |
| 1. | Bulk density | 1.52 g/cm<sup>3</sup> | Core sampling (Blake and Hartge, 1986) |
| 2. | pH and | 7.41 | 1:2.5 (w/v) soil-water suspension (Kumar et al. |
| 3. | Electrical Conductivity | 0.24 dS/m | 2011) |
| 4. | Oxidizable OC | 7.80g/kg | Wet digestion (Walkley & Black, 1934) |
| 5. | Available N | 222 kg/ha | Alkaline permanganate (Subbiah & Asija, 1956) |
| 6. | Available P | 25 kg/ha | Olsen method (Olsen S.R.1954) |
| 7. | Available K | 297 kg/ha | Ammonium acetate (Hanway and Heidel, 1952) |

### 2.2 Experimentation

The experiment was conducted with three rice-based cropping systems, namely, RMCp, rice-RWGg and RCfBr following split plot design (SPD). Tillage intensity CT, MT, and ZT, was main plot, while R_0_ (0% Residue and 100% RDF), R_50_ (50% Residue and 100% RDF & 50% Residue and 75% RDF) and R_100_ (100% Residue and 50% RDF &100%Residue and 75%RDF) was sub-plot treatment. Each plot was replicated thrice in plot of size 1720m^2^. Details of the crop management practices and treatments are described in Fig 1 & Table 2a.

**Fig 1:**
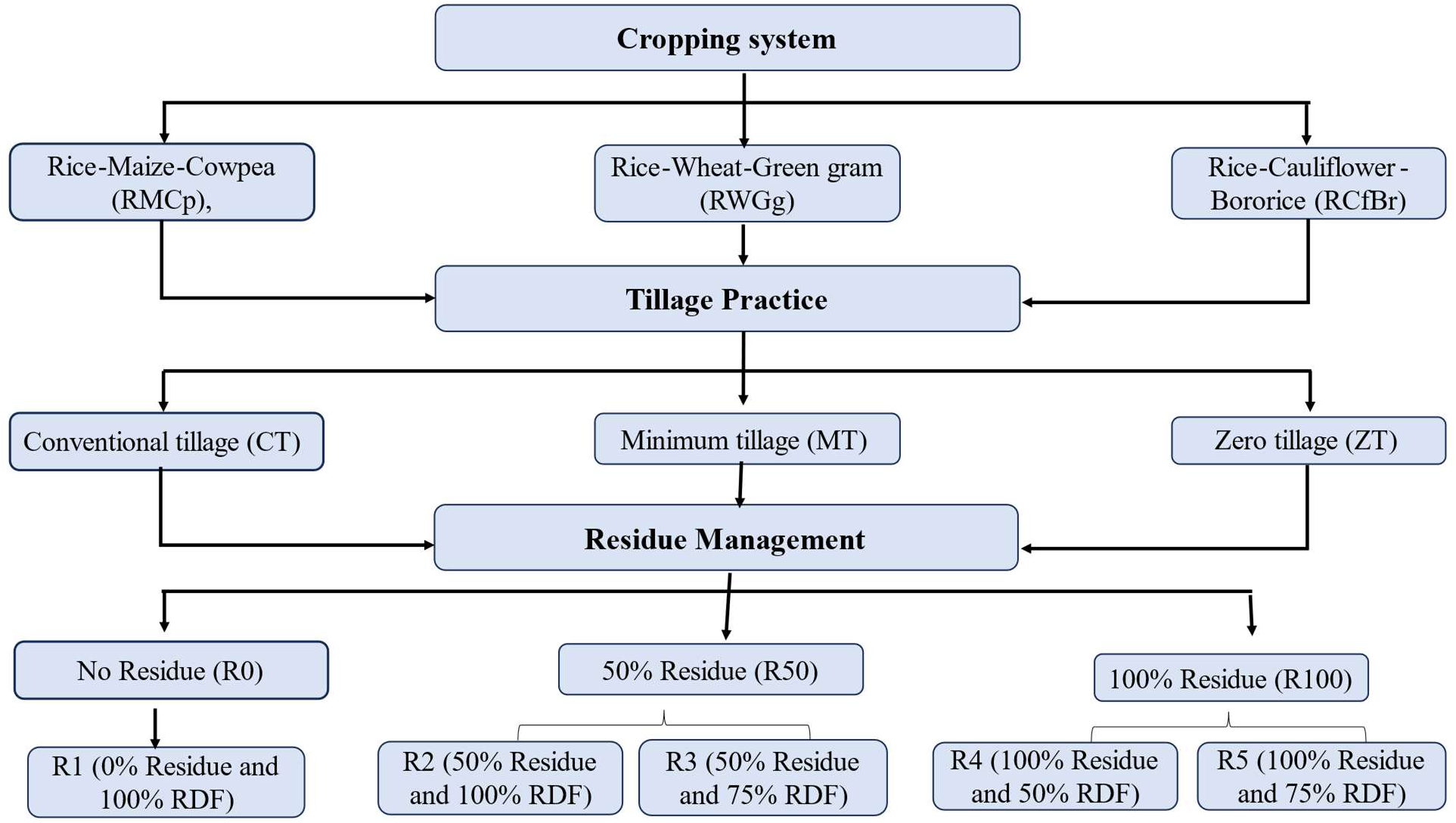
Cropping system, tillage intensity and residue management used in experiment

**Table 2a:** Crop management practices used for used for crop selected for experimentation.

| Crop management practices | RMCp |  |  | RWGg |  |  | RCfBr |  |  |
| --- | --- | --- | --- | --- | --- | --- | --- | --- | --- |
|  | Rice | Maize | Cowpea | Rice | Wheat | Green gram | Rice | Cauliflower | Boro rice |
| Sowing | In ZT sowing was done using zero-till-seed-drill, while in MT and CT plots by multi-crop seed-cum-fertilizer-drill was used. |  |  |  |  |  |  |  |  |
| Fertilizer management | Rice | Maize | Cowpea | Rice | Wheat | Green gram | Rice | Cauliflower | Boro rice |
| N: P<sub>2</sub>O<sub>5</sub>: K<sub>2</sub>O (Kg/ha) | 80:40:40 | 140:70:70 | 20:60:60 | 80:40:40 | 120:60:60 | 20:40:20 | 80:40:40 | 180:90:90 | 120:60:60 |
| Fertilizer application | Time of application: as basal; Fertilizer used for supplying nutrients: Urea, and customized 10.0- 11.4-21.7 (N-P-K) fertilizer |  |  |  |  |  |  |  |  |
| Residue load* (Kg/ha) | 2018 | 2019 | 2020 | 2018 | 2019 | 2020 | 2018 | 2019 | 2020 |
|  | 0% = 0 | 0% = 0 | 0% = 0 | 0% = 0 | 0% = 0 | 0% = 0 | 0% = 0 | 0% = 0 | 0% = 0 |
|  | 50%=2270 | 50%= 3092 | 50%= 2985 | 50%= 2270 | 50%= 3092 | 50%= 3285 | 50%= 2270 | 50%= 2590 | 50%= 3210 |
|  | 100%=4540 | 100%=6185 | 100%= 5969 | 100%=4540 | 100%=6185 | 100%=6571 | 100%=4540 | 100%=5181 | 100%= 6420 |
| Irrigation | Irrigation was done at 50% moisture deficit in soil to maintain sufficient moisture for crops. In rice total number of irrigations ranged from 6 to 8 depending upon the amount and distribution of rainfall in different years. In each irrigation, 5-6 cm of water was applied. |  |  |  |  |  |  |  |  |
| Intercultural operation | Weeds were controlled by application of pre-emergence and post-emergence herbicides as and when required. In addition, glyphosate (41% SL) was applied, especially for ZT @ 750 g a.i. ha<sup>-1</sup> (7-10 days before sowing). |  |  |  |  |  |  |  |  |

151 \*Rice residue was retained for the succeeding *Rabi* crops

### 2.3 Soil Sampling and Analysis

Soil samples were collected on completion of each cropping cycle (in pre-*kharif of* 2019 & 2020) from the rhizosphere. From each plot, 5–6 plants were uprooted and soil adhering to their root were extracted and mixed properly to make composite sample before transferring to plastic bag and stored in refrigerator at 4°C. Soil enzymes were estimated by methods given in Table 2b. Microbial activities were measured by incubating soil samples in 2-set of conical flasks containing broth (before & after sterilization) for required period. SMB (SMBC and SMBN) was estimated by chloroform-fumigation extraction (CFE) method, followed by incubation at 25°C for 10 days of fumigated and unfumigated soil samples, with soil moisture < 55% of water-holding capacity.

**Table 2b:**
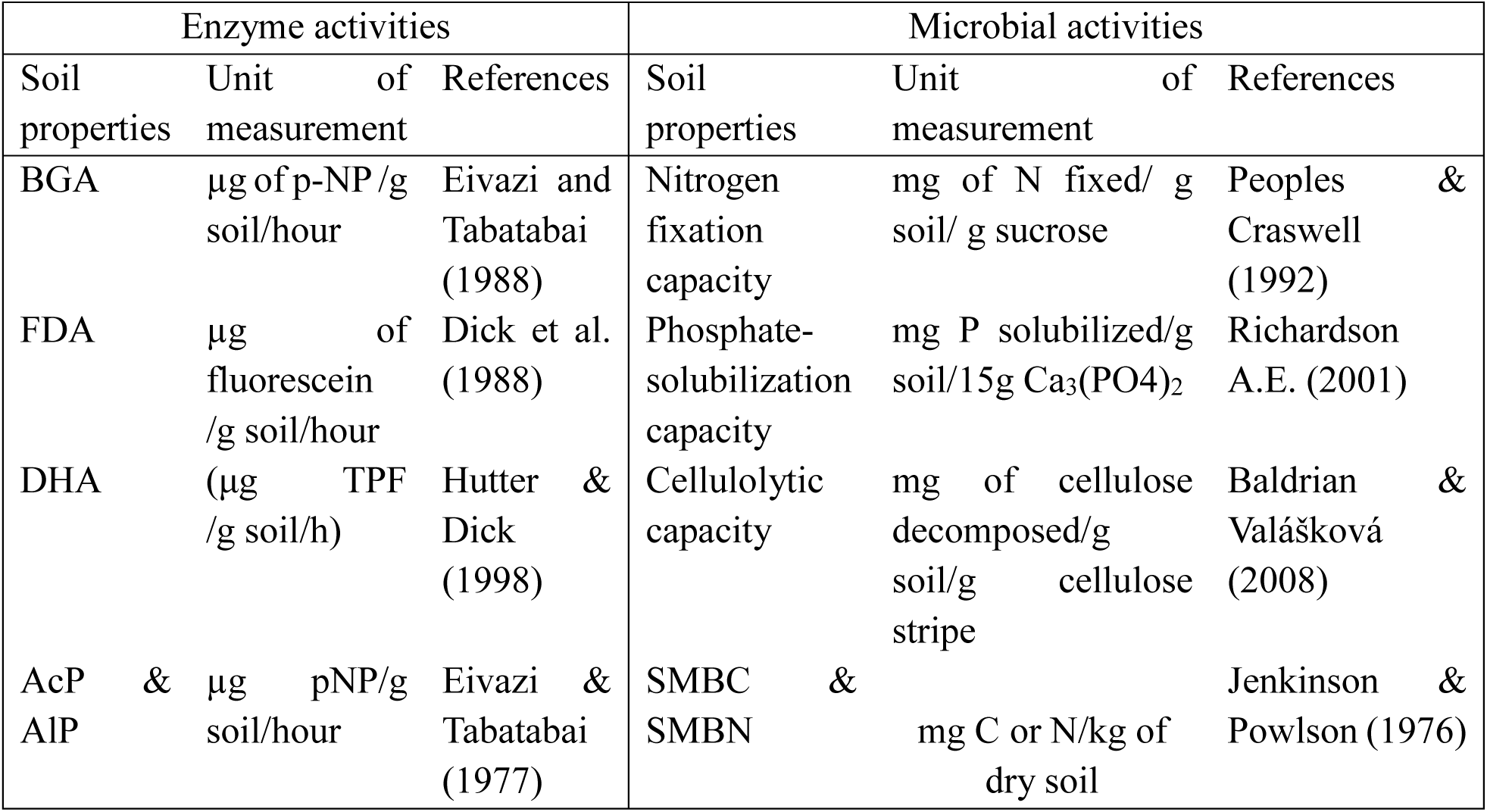
Methods used for the analysis of different biochemical and microbial properties of soil.

### 2.4 Statistical Analysis

Microbial and biochemical properties of soil under CA were subjected to analysis of variance (ANOVA) using the generalized linear model on SPD to determine impact of tillage, residue & nutrient, cropping system and their interactions (fixed effects). Differences among treatemnts were compared at 5% probability level through Duncan’s multiple range test (DMRT) using Statistical Package for the Social Sciences (SPSS) software (version 20.0). Means for treatment effects were separated based on the least significant difference (LSD). The LSD values were tested at (*P < .05*) probability level.

### 2.5 Suitability of conservation practices with respect to biochemical & microbial properties of soil

To identify suitability of CA with respect to tillage practices, residue level and cropping system for sustaining soil biochemical and microbial properties, a multivariate assessment (Principal component analysis, PCA) was done using variables DHA, BGA, FDA, AcP, AlP, NFBAct, PSBAct, CDBAct, Bacteria, Fungi, Actinomycetes, CDB, PSB, NFB, AMF, SOC, Available N, Available P, SMBC and SMBN. Two principal components (PC1 and PC2) with eigen value >1 were chosen. Based on factor loading of both the PCs total score was calculated. Overall, suitability of conservation practices was examined from the magnitude of total score in ascending order (Mukherjee et al. 2024). The weighted additive indexing approach was utilized to carry out indicator selection, interpretation, and integration into the overall BSHI employing. The variables that best describe the system characteristics were believed to be the main components with high eigen values and factor loadings (Brejeda et al. 2000) and included in minimum data set (MDS). The score functions for each variable were converted or normalised to a value between 0 (least favourable soil function) and 1 (most favourable soil function) (Andrews et al. 2002). To calculate the biological soil health index (BSHI), the MDS variables for each observation were weighted using the PCA scoring. Weighted additive BSHI was calculated by using the following formula:

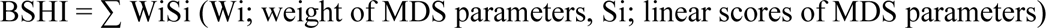

## 3. Results

### 3.1 Impact of tillage intensity and residue application on soil enzymes under different cropping system

A dynamic change in enzyme activity was observed throughout the course of experimentation. Activities of different enzymes ranged from 14.07±2.48-21.62±1.37 µg TPF/g-soil/h, 78.96±1.24-91.70±1.34 (µg pNP/g-soil/ h), 71.23±2.19-75.90±1.51(µg pNP/g-soil/ h), 7.71±0.59 -11.44 ±0.52 (µg pNP/g-soil/ h), and 16.99±1.88 to 18.75±1.67 µg fluorescein/g-soil/ h for DHA, AlP, AcP & FDA, respectively. DHA, BGA and AlP were highest in ZT (Parihar et al. 2016) while FDA and AcP showed non-significant effect of tillage practices among cropping system studied. AlP showed varied response of different tillage intensities among cropping system under study (Fig.2 (i), (ii) & (iii)). Under RMCp, R_50_ & R_100_ excelled over R_0_ (Fig. 3(i)), while R_100_ performed was significantly higher compared to R_50_ and R_0_ under RWGg & RCfBr particularly for BGA and AlP. However, enzymes AcP and FDA showed non-significant influence of residue application (Fig. 3 (i) & (ii)). Lower enzymatic activities were seen under different cropping system with higher RDF (Dick et al. 1997). Treatment R_4_ (100% residue and 50% RDF) excelled under RWGg & RCfBr (Liang et al. 2014). DHA, BGA & AlP excelled under RMCp & RWGg, however, activities of AcP & FDA was at par activities under different cropping system (Fig. 4).

**Fig. 3:**
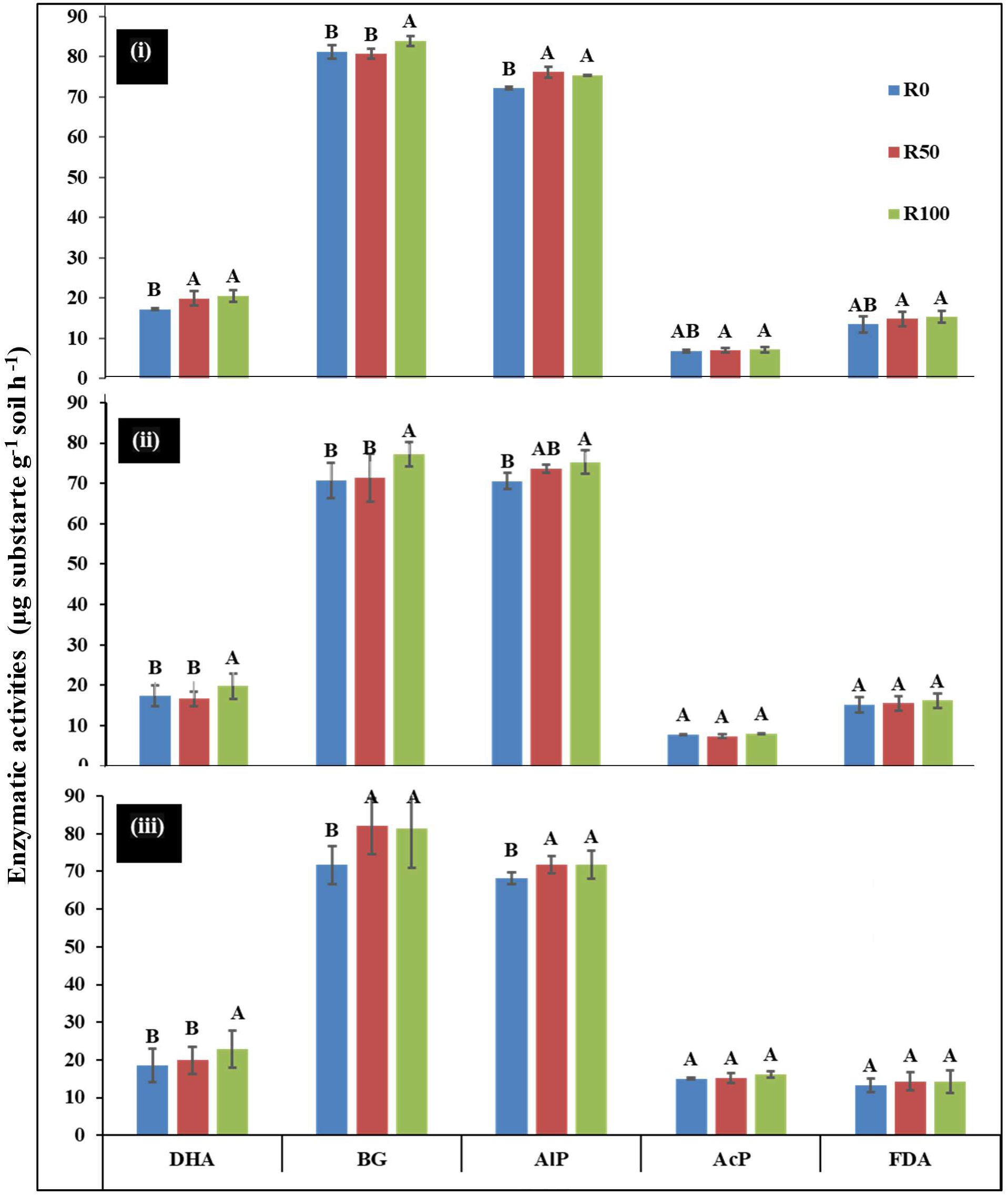
Enzymatic activities in soil under different level of residue application in (i) Rice-Maize-Cowpea, (ii) Rice-Wheat-Green gram & (iii) Rice-Cauliflower-Bororice/summer rice cropping system (R_0_: 0% Residue and 100% RDF, R_50_: 50% Residue and 100% RDF & 50% Residue and 75% RDF and R_100_:100% Residue and 50% RDF & 100% Residue and 75% RDF; Error bars indicate (±) SEm of the observed values. Values with capital alphabets (A, B) indicates significant difference among doses of residue application at 5% level)

**Fig.2:**
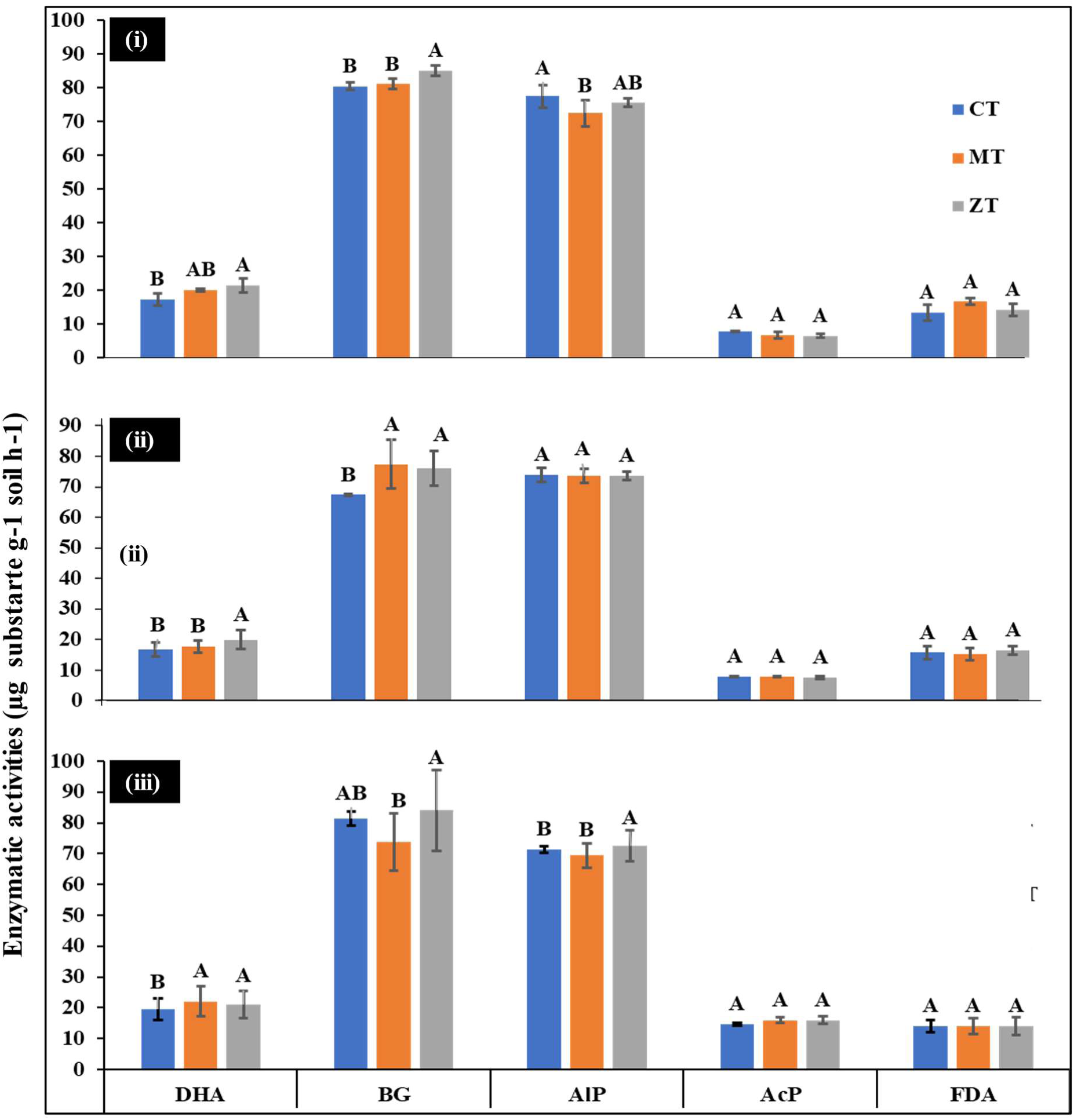
Enzymatic activities in soil under different tillage practices applied under (i) Rice-Maize-Cowpea, (ii) Rice-Wheat-Green gram & (iii) Rice-Cauliflower-Bororice/summer rice cropping system (CT= Conventional tillage, MT= Minimum tillage and ZT= Zero tillage; Substrate for DHA is TPF (Triphenyl formazone), for BGA, AlP & AcP is pNP (Para nitrophenol), & for FDA it is fluorescein diacetate; error bars indicate (±) SEm of the observed values. Values with capital alphabets (A, B) indicates significant difference among tillage practices at 5% level)

**Fig 4:**
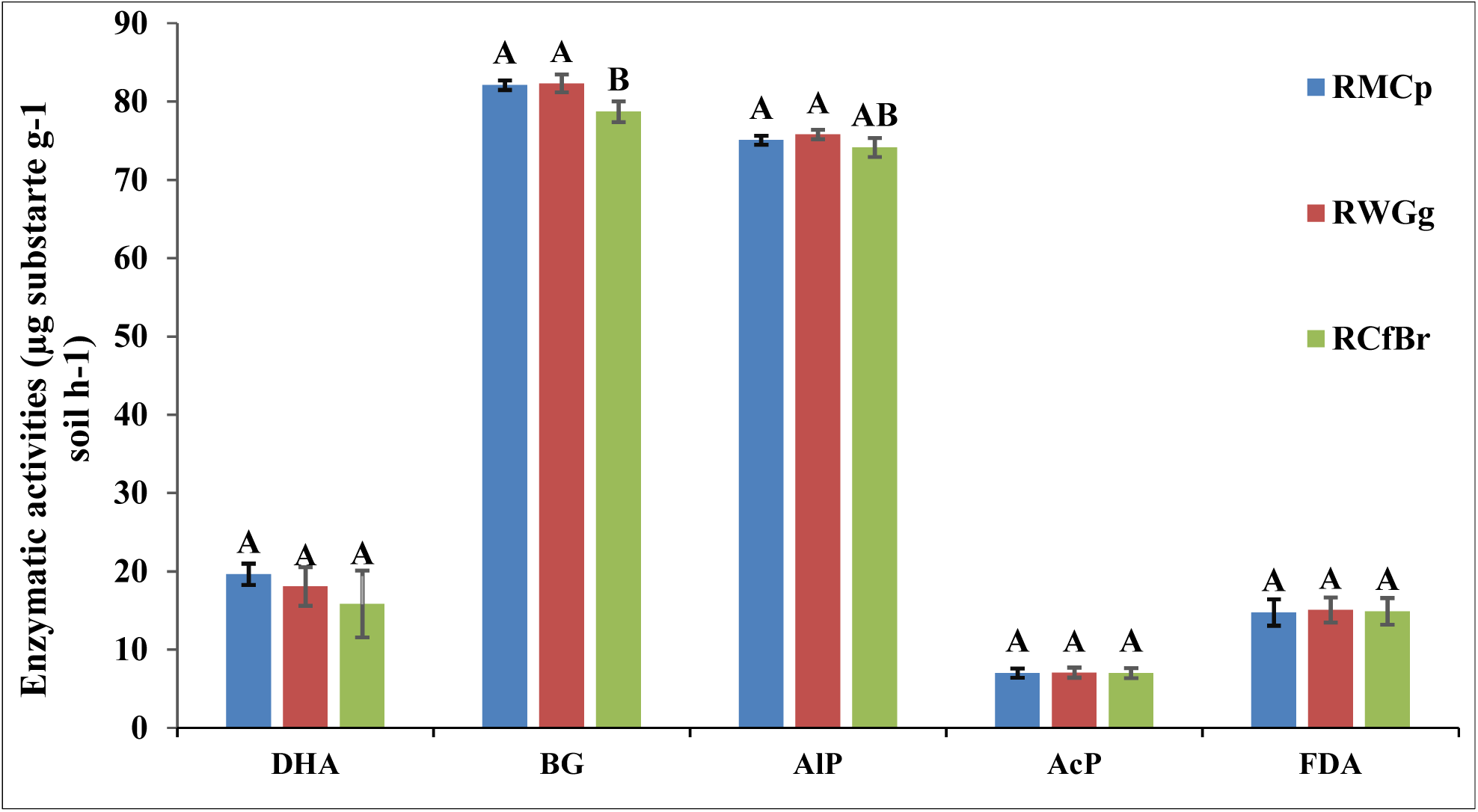
Soil enzyme activities under different cropping system following CA (RMCp: rice-maize-cowpea, RWGg: rice-wheat-green gram and RCfBr: Rice-cauliflower-bororice/summer rice; Error bars indicate (±) SEm of the observed values. Values with capital alphabets (A, B) are indicates significant difference among cropping systems at 5% level)

### 3.2 Impact of tillage intensity and residue application on soil microbial activities under different cropping system

Low input-farming e.g., CA is designed to rejuvenates soil microbes, thus activities of selected microbes (FLNF, PSB and cellulolytic microbes) was studied. NFBAct & PSBAct followed the pattern as observed in case of soil enzymes, though CDBAct showed reverse pattern wrt tillage and residue application. NFBAct ranged from 0.29 ± 0.01 to 0.41 ± 0.01 mg-N fixed/g-soil/g-sucrose under CA, while PSBAct and CDBAct ranged from 0.64±0.06 to 1.34±0.27mg-P/g-soil/15g Ca_3_(PO_4_)_2_ and 83.15±10.11 to 138.89±16.64 mg-cellulose decomposed/g-soil /g-cellulose stripe, respectively. A gradual increase in microbial activities with increase in level of residue application was observed. R_100_ showed significantly higher NFBAct & PSBAct than R_0_ and R_50_. Among cropping system, RMCp and RWGg (legume-based) excelled over RCfBr (cereal-based). Combining the effect of tillage and residue, R_100_ under MT and ZT was more effective, except for CDBAct (Fig. 5 (i), (ii) & (iii).). In RCfBr, CDBAct, & NFBAct reported to be highest in CT with R_100_ (Fig. 5(iii)).

**Fig 5:**
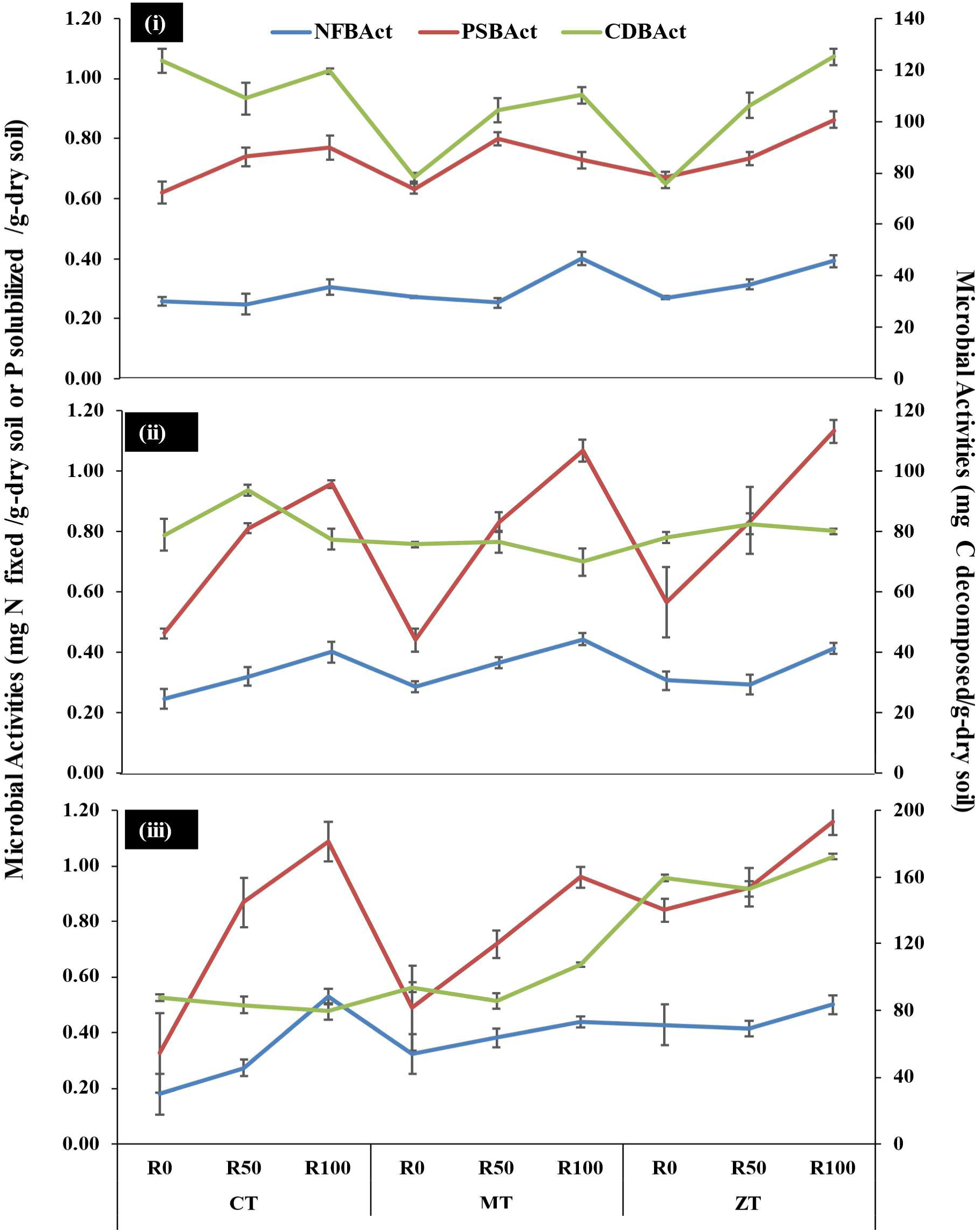
Microbial activities {Nitrogen fixation (NFBAct), Phosphate solubilization (PSBAct) & Cellulolytic activities (CDBAct) under different tillage and residue management in (ii) Rice-Maize-Cowpea, (ii) Rice-Wheat-Green gram & (iii) Rice-Cauliflower-Bororice/summer rice cropping system, where CT= Conventional tillage, MT= Minimum tillage and ZT= Zero tillage, R_0_: 0% Residue and 100% RDF, R_50_: 50% Residue and 100% RDF & 50% Residue and 75% RDF and R_100_:100% Residue and 50% RDF & 100% Residue and 75% RDF. Error bars indicate (±) SEm of the observed values

### 3.3 Effect of tillage and residue management on soil microbial population under different cropping system

Bacterial population was at par with different degree of tillage intensities indicating bacteria are not effectuated of soil disturbance under different tillage practices. In case of fungi and actinomycetes degree of soil disturbance was negatively associated with their population i.e., ZT harboured maximum number while CT retained least population in soil (Fig 6 (a), (b) & (c)). Among specific microbial groups like FLNF, PSB and CDB, population varied from 37±1.87 to 42±1.77 CFUx10^6^, 32±1.33 to 39±1.59 CFUx10^5^ and 40±1.43 to 47±1.15 CFUx10^3,^ respectively among tillage practices. R_100_ restored highest microbial population over R_50 &_ R_0_. Likewise, soil enzymes and microbial activities, broad microbial group also showed better performance under legume-based cropping system (RMCp and RWGg) compared to RCfBr. Higher doses of residue application were more effective under CT than MT and ZT as indicated in RMCp. However, PSB and FLNF showed higher response towards higher residue application irrespective of tillage intensities (Fig. 6a). Under RWGg, conservation tillage appeared favorable for restoring fungi and actinomycetes (hyphal microbes), while specific soil microbes showed increasing trend with higher residue application (Fig. 6b). Under RCfBr, different microbial group responded differently. Bacteria and fungi population showed gradual increase in population with minimizing soil disturbance. However, actinomycetes were highest in ZT (Fig. 6c). Specific soil microbes showed higher population towards combine effect of CA practices (tillage and residue application).

**Fig. 6a:**
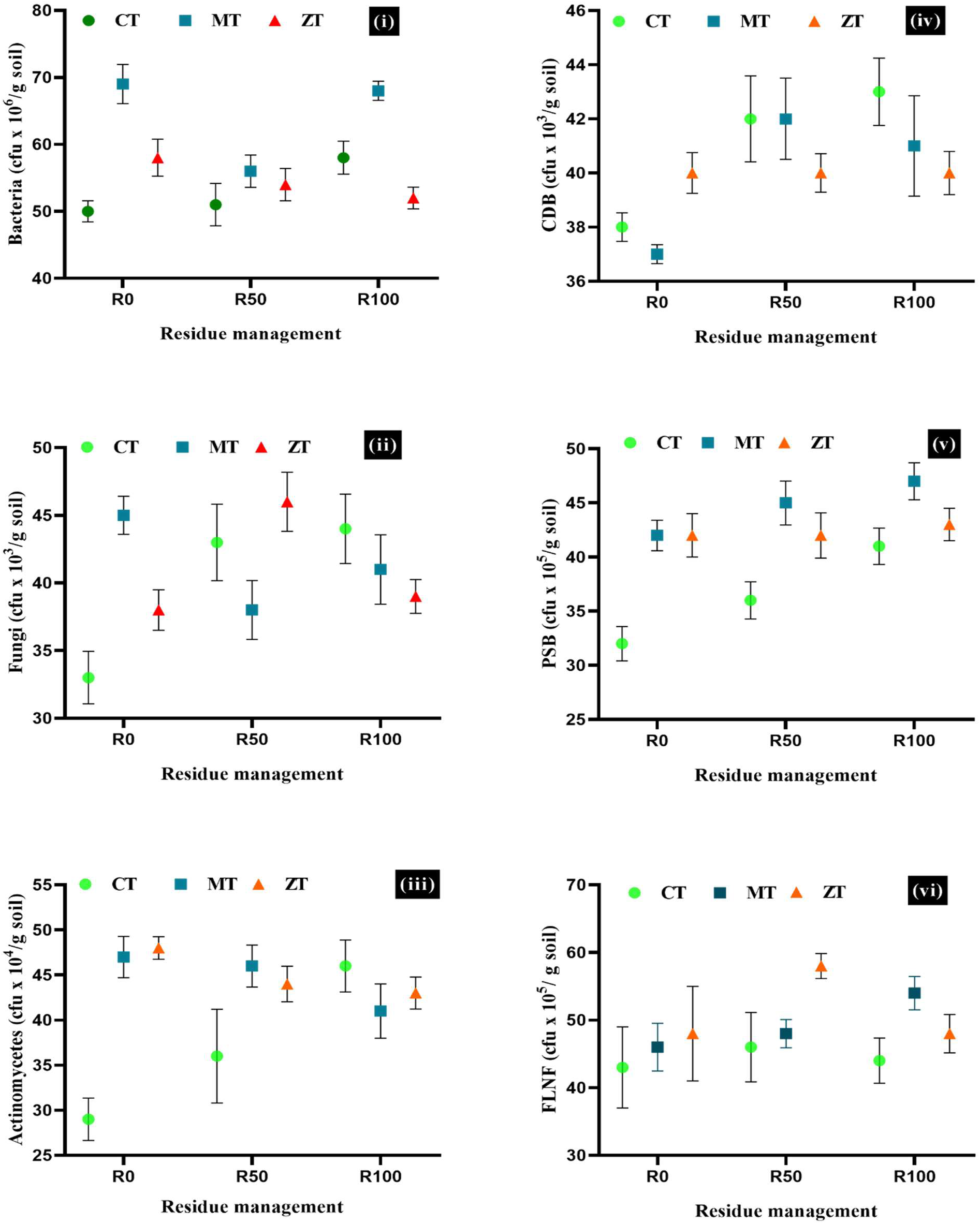
Population of specific microbial groups in soil under different tillage and residue management in Rice-Maize-Cowpea cropping system, where where CT= Conventional tillage, MT= Minimum tillage and ZT= Zero tillage, R_0_: 0% Residue and 100% RDF, R_50_: 50% Residue and 100% RDF & 50% Residue and 75% RDF and R_100_:100% Residue and 50% RDF & 100% Residue and 75% RDF. Error bars indicate (±) SEm of the observed values

**Fig 6b:**
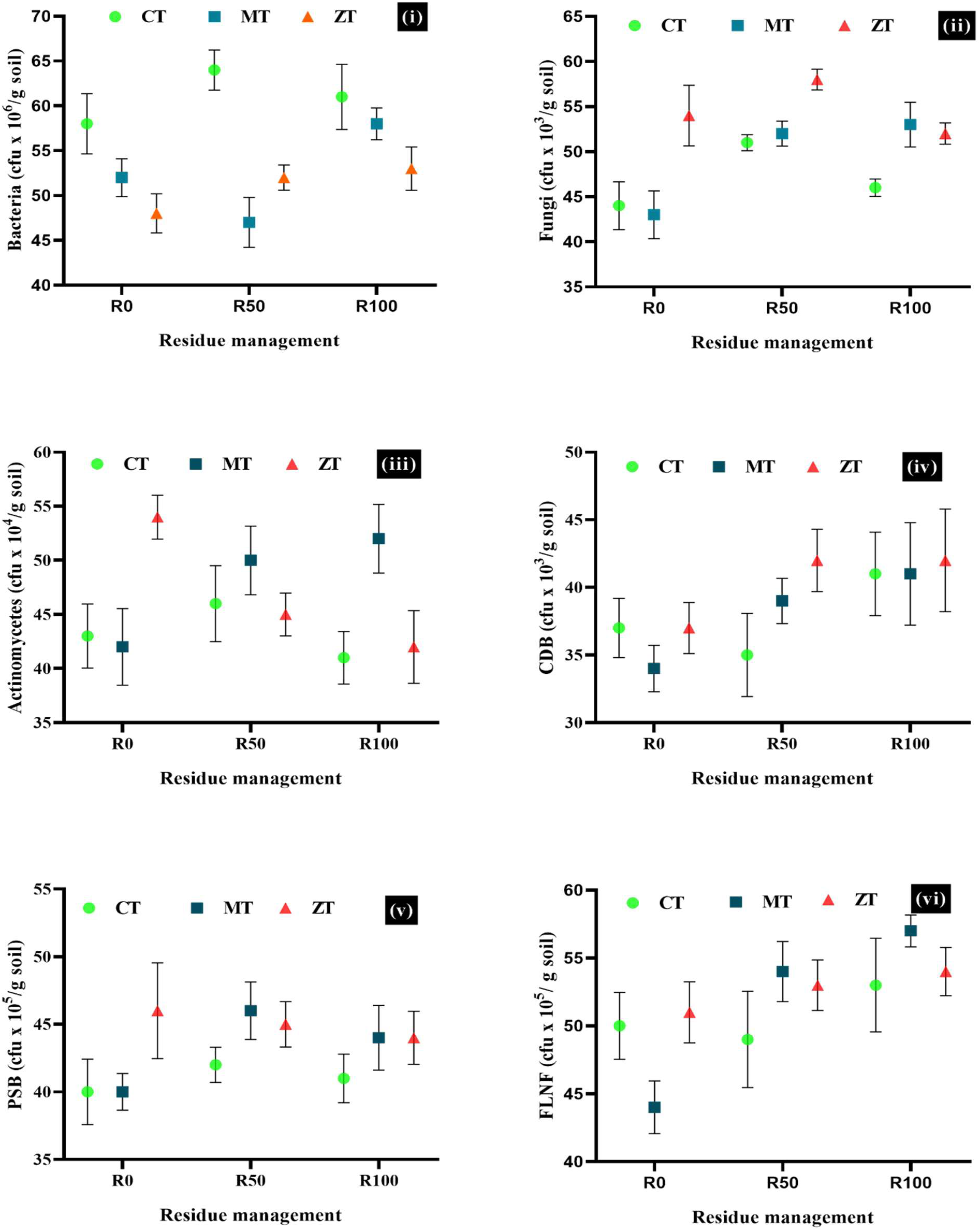
Population of specific microbial groups in soil under different tillage and residue management in Rice-Wheat-Green gram cropping system, where where CT= Conventional tillage, MT= Minimum tillage and ZT= Zero tillage, R_0_: 0% Residue and 100% RDF, R_50_: 50% Residue and 100% RDF & 50% Residue and 75% RDF and R_100_:100% Residue and 50% RDF & 100% Residue and 75% RDF. Error bars indicate (±) SEm of the observed values

**Fig 6c:**
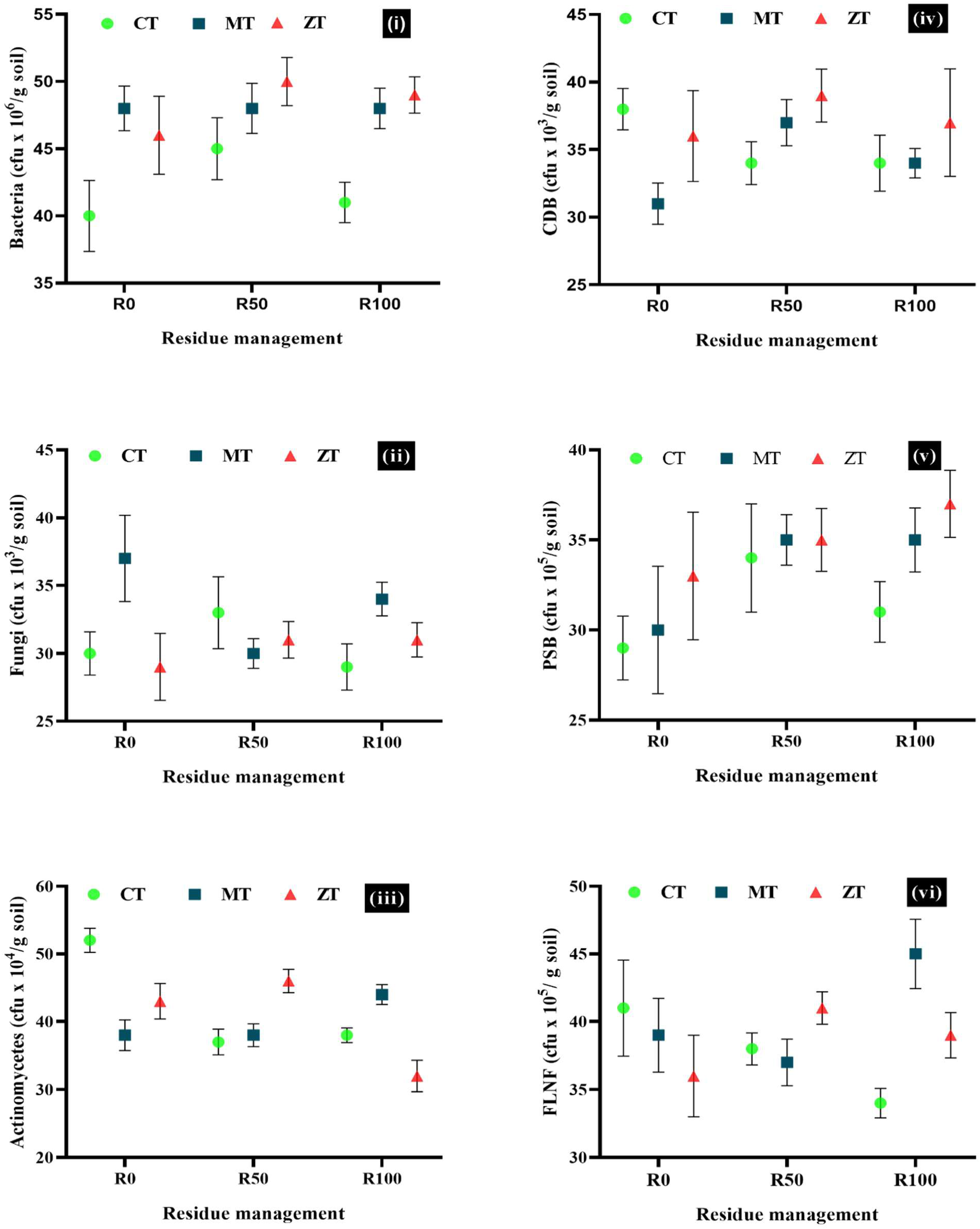
Population of specific microbial groups in soil under different tillage and residue management in Rice-Cauliflower-Bororice/summer rice cropping system, where CT= Conventional tillage, MT= Minimum tillage and ZT= Zero tillage, R_0_: 0% Residue and 100% RDF, R_50_: 50% Residue and 100% RDF & 50% Residue and 75% RDF and R_100_:100% Residue and 50% RDF & 100% Residue and 75% RDF. Error bars indicate (±) SEm of the observed values

### 3.4 Characterization of chemical properties and microbial biomass in soil with different level of tillage intensity and residue application under different cropping system

Chemical properties studied were at par under MT & ZT with respect to available N and SOC while in case of available P CT was at par to ZT. Among different tillage practices available N, SOC and available P varied from 258.66±2.53 to 267.54±2.57 kg/ha, 9.27±0.76 to 9.99±0.58 g/kg-soil & 31.79±2.25 to 34.06±2.75 kg/ha, respectively. Similarly, under residue treatment they varied from 260.02±1.97 to 266.50±1.93 kg/ha, 9.04±0.07 to 10.09±0.06 g/kg- soil & 31.88±1.91 to 35.83±3.83 kg/ha while under cropping system it ranged from 261.37±3.29 to 266.34±1.78 kg/ha, 9.47±0.26 to 10.39±0.53 g/kg-soil & 32.06±1.54 to 33.33±2.78 kg/ha, respectively (Fig 7a & 7b). Residue retention increased SOC, available N & P in soil (Bhattacharyya et al. 2019). SMBC and SMBN was higher under ZT (254.07±2.70 mg C/kg-dry soil & 49.73±2.05 mg N/kg- dry soil, respectively) compared to CT (238.47±2.71 mg C/kg-dry soil & 46.67±2.49 mg N/kg-dry soil, respectively) (Zuber and Villamil, 2016). SMBC & SMBN varied from 237.95±5.55 to 251.05±5.65 mg C/kg-dry soil & 44.66±2.51 to 49.37±1.26 mg N/kg-dry soil, respectively under different level of residue application (Fig 7c). Under RWGg and RCfBr, tillage intensity showed negative response with SOC, however this trend was different in RMCp, where CT outperformed conservation tillage (Fig 7a). Gradual increase in SOC was observed with doses of residue application.

**Fig 7a:**
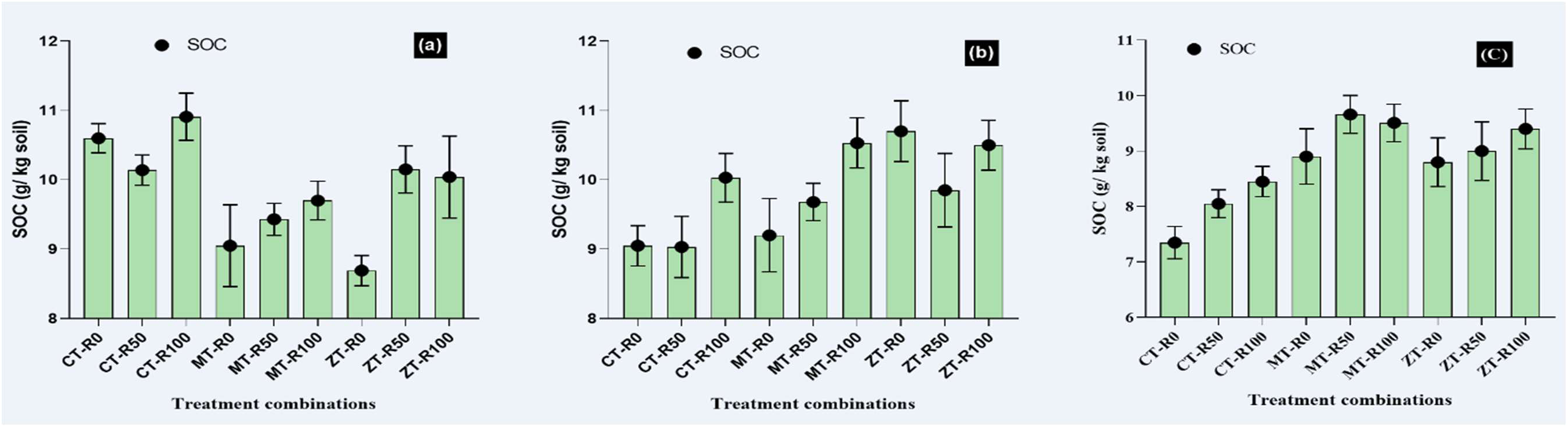
Impact of CA on SOC under (a) rice-maize-cowpea (RMCp); (b) rice-wheat-green gram (RWGg) & (c); rice-cauliflower-bororice/summer rice (RCfBr) cropping system, where where CT= Conventional tillage, MT= Minimum tillage and ZT= Zero tillage, R_0_: 0% Residue and 100% RDF, R_50_: 50% Residue and 100% RDF & 50% Residue and 75% RDF and R_100_:100% Residue and 50% RDF & 100% Residue and 75% RDF. Error bars indicate (±) SEm of the observed values.

**Fig 7b:**
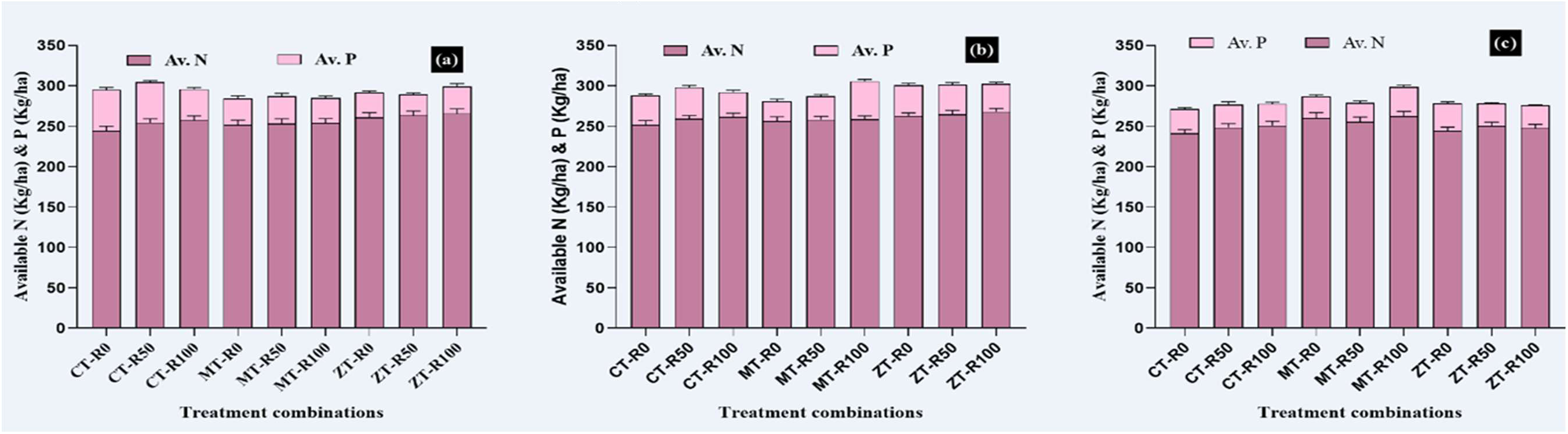
Impact of CA on available phosphorus and nitrogen under (a) rice-maize-cowpea (RMCp); (b) rice-wheat-green gram (RWGg) & (c); rice-cauliflower-bororice/summer rice (RCfBr) cropping system, where where CT= Conventional tillage, MT= Minimum tillage and ZT= Zero tillage, R_0_: 0% Residue and 100% RDF, R_50_: 50% Residue and 100% RDF & 50% Residue and 75% RDF and R_100_:100% Residue and 50% RDF & 100% Residue and 75% RDF. Error bars indicate (±) SEm of the observed values.

**Fig 7c:**
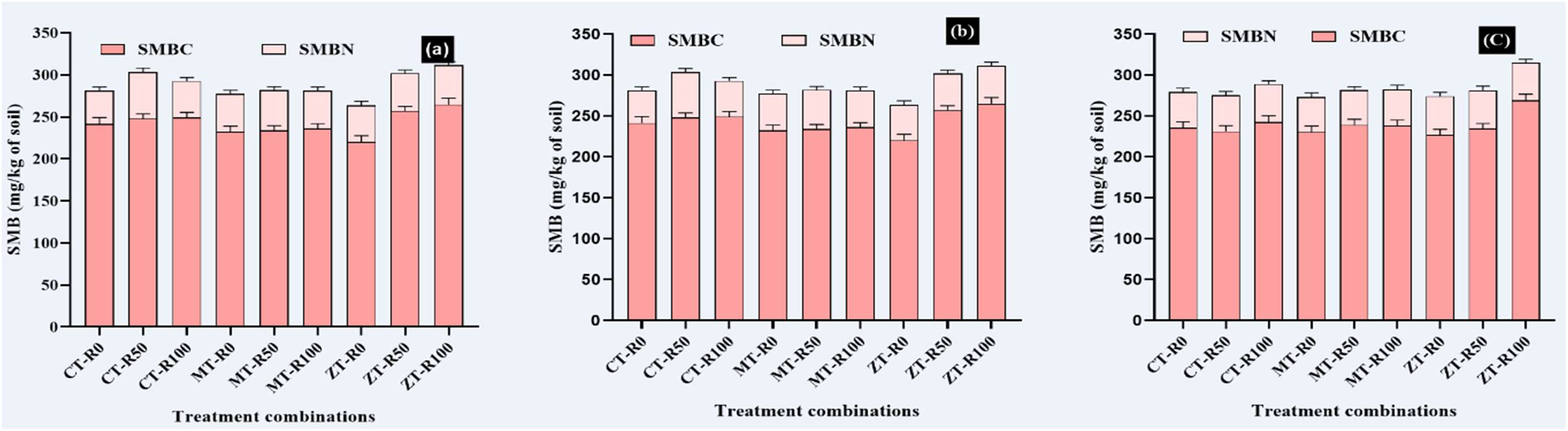
Impact of CA on soil microbial biomass carbon (SMBC) and soil microbial biomass nitrogen (SMBN) under (a) rice-maize-cowpea (RMCp); (b) rice-wheat-green gram (RWGg) & (c); rice-cauliflower-bororice/summer rice (RCfBr) cropping system, where where CT= Conventional tillage, MT= Minimum tillage and ZT= Zero tillage, R_0_: 0% Residue and 100% RDF, R_50_: 50% Residue and 100% RDF & 50% Residue and 75% RDF and R_100_:100% Residue and 50% RDF & 100% Residue and 75% RDF. Error bars indicate (±) SEm of the observed values.

### 3.5 Impact of CA on biochemical, microbial and chemical properties of soil based on ANOVA (Analysis of Variance)

ANOVA showed non-significant influence of tillage on DHA while significant effect on BGA, AlP, AcP and FDA (p≤0.05). CS x RM had significant effect on DHA, BGA, AlP, AcP and FDA (Table 3). ANOVA showed highly significant difference among main source of variation, (T, CS and RM) and their interaction on bacteria, fungi and actinomycetes count. Bacteria showed non-significant variation towards T, while highly significant variation toward CS and RM (p ≤ 0.01) and interaction between T × CS, RM × CS, RM × T and T × RM × CS (Table 4). Fungi and actinomycetes showed highly significant difference towards T and CS (p≤0.01) and RM (p≤0.01), interaction between T × CS, RM × T, RM × CS and RM ×T × CS. ANOVA technique revealed non-significant variation of T on CDB, significant variation (p≤ 0.05) on PSB while highly significant (p≤0.01) on FLNF. Towards CS and RM, specific microbial group showed highly significant variation (p≤0.01). Interaction of T × CS on FLNF and CDB showed significant variation (p≤0.05) while PSB showed highly significant variation (p≤0.01). Interaction between RM × T, RM × CS, RM× T ×CS, specific microbial groups (FLNF, PSB, CDB) showed highly significant variation. RM×CS showed significant variation at p≤0.05 in case of PSB.

**Table 3:** Impact of CA on soil enzymes based on ANOVA.

| Source of variance (SV) | df | Mean square |  |  |  |  |
| --- | --- | --- | --- | --- | --- | --- |
|  |  | DHA | BGA | AIP | AcP | FDA |
| Tillage (T) | 2 | 68.734ns | 59.05* | 59.73* | 0.29* | 7.040* |
| Error a | 2 | 6.301 | 1.00 | 0.19 | 0.00 | 0.022 |
| Cropping system (CS) | 2 | 57.513** | 577.97** | 137.50** | 758.73** | 23.729** |
| T x CS | 4 | 9.106 | 301.06** | 20.51** | 5.35** | 14.023** |
| Error b | 6 | 3.672 | 1.70 | 0.19 | 0.82 | 0.056 |
| RM (Residue Management) | 4 | 41.966** | 154.84** | 121.90** | 2.17xx | 10.055** |
| RM x T | 8 | 5.387* | 59.36** | 65.64** | 1.23* | 2.425** |
| RM x CS | 8 | 7.484** | 77.53** | 6.71** | 1.01** | 1.801** |
| RM x T x CS | 16 | 9.589** | 31.37** | 118.35** | 2.19** | 1.974* |
| Error c | 36 | 2.297 | 7.19 | 0.26 | 0.01 | 0.009 |
| CV (a) % |  | 12.9 | 1.3 | 0.6 % | 0.3 % | 0.9 % |
| CV (b) % |  | 9.8 | 1.7 | 0.6 | 8.5 % | 1.5 % |
| CV (c) % |  | 7.8 | 3.4 | 0.7 | 0.7 % | 0.6 % |
| Mean |  | 19.51 | 78.466 | 77.68 | 10.70 | 15.73 |

(ns: non-significant; ‘\*\*\*’ Significant at the 0.01 probability level; ‘\*\*’ Significant at the 0.05 probability level; df: degrees of freedom; CV: coefficient of variation)

**Table 4:** Impact of CA on microbial population and their activities based on ANOVA.

| Source of variance (SV) | df | Mean square |  |  |  |  |  |  |  |  |
| --- | --- | --- | --- | --- | --- | --- | --- | --- | --- | --- |
|  |  | Bacteria | Fungi | Actinomycetes | FLNF | PSB | CDB | Nitrogen fixation | Phosphate solubilization | Cellulolytic activities |
| Tillage (T) | 2 | 88.7ns | 37.73** | 169.211** | 336.04** | 18.98ns | 14.144ns | 0.025574 ns | 0.10749ns | 6100.3** |
| Error a | 2 | 5.7 | 0.04 | 0.223 | 3.38 | 3.73 | 6.144 | 0.001990 | 0.00964 | 17.5 |
| Cropping system (CS) | 2 | 923.1** | 320.53** | 240.544** | 1313.41** | 845.21** | 232.578** | 0.063481** | 0.13234** | 10067.3** |
| T × CS | 4 | 180.7** | 44.47* | 28.344* | 21.46* | 29.28** | 27.178* | 0.007401** | 0.03253** | 6408.3** |
| Error b | 6 | 11.0 | 6.44 | 4.322 | 4.09 | 2.46 | 5.000 | 0.001563 | 0.00102 | 22.2 |
| RM (Residue Management) | 4 | 55.3** | 54.53** | 166.806** | 210.60** | 71.96** | 94.517** | 0.001563** | 0.52521** | 397.7** |
| RM × T | 8 | 45.6** | 41.26** | 39.114** | 39.06** | 13.06** | 33.408** | 0.002123** | 0.01285** | 283.7 ** |
| RM × CS | 8 | 49.3** | 25.19** | 55.947** | 19.34** | 9.13* | 43.842** | 0.003213** | 0.05615** | 392.7 ** |
| RM × T × CS | 16 | 25* | 18.24** | 101.206** | 16.23** | 16.53** | 22.608** | 0.006039** | 0.01100** | 211.2 ** |
| Error c | 36 | 12.6 | 4.09 | 6.917 | 3.17 | 3.32 | 6.139 | 0.000474 | 0.00141 | 27.2 |
| CV (a) % |  | 5.3 | 0.6 | 1.1 | 3.9 | 4.9 | 6.5 | 12.6 | 11.9 | 4.2 |
| CV (b) % |  | 7.3 | 6.8 | 4.8 | 4.3 | 4 | 5.85 | 11.1 | 3.9 | 4.7 |
| CV (c) % |  | 7.9 | 5.4 | 6.1 | 3.8 | 4.6 | 6.4 | 6.1 | 4.6 | 5.2 |
| Mean |  | 45.288 | 37.5 | 42.88 | 47.42 | 39.19 | 38.42 | 0.3547778 | 0.823 | 100.43 |

(ns: non-significant; \*\*\* Significant at the 0.01 probability level; \*\* Significant at the 0.05 probability level; df: degrees of freedom; CV:
coefficient of variation)

### 3.6 Correlation of soil enzyme and microbial activities with microbial and chemical properties of soil

DHA showed a significant positive correlation with fungi, actinomycetes, FLNF, PSB, CDB (except bacteria), SOC and available N (Jha et al. 1992) in soil under CA, while BGA with soil microbes e.g., FLNF (r= 0.593, p≤ 0.05) and CDB (r=0.537, p≤ 0.05) as well as SOC (r= 0.815, p ≤ 0.01) & SMBC (r= 0.516, p ≤ 0.05). Phosphatase activity served as good soil quality indicator because of its strong correlation with SOC, available P, and N. However, AcP showed positive correlation (r= 0.592, p≤ 0.05) with available N while same with AlP showed negative correlation (r= -0.605, p≤ 0.05) (Adetunji et al. 2017). FDA showed highly significant correlation with PSB, FLNF, AMF, SMBC, SOC, available N and P in soil. DHA is strongly correlated with fungal and actinomycetes population than other soil enzymes. Available N was significantly correlated with all the soil enzymes studied except BGA (Table 6). Out of five enzymes studied, DHA and FDA were significant and positively correlated with most of the biochemical parameter evaluated. CDB was found to be significantly correlated with NFBAct, which showed significant positive correlation with PSB, available P and SOC in soil, however activity was significantly (negatively) correlated with soil available N. PSBAct showed significant correlation with soil microbes (e.g., Fungi, FLNF, PSB, AMF etc.), SMBC, available N as well as SOC (Table 6). However, PSBAct as well as population of PSB are found to be strongly negatively associated with soil available P (Zheng et al. 2021). Cellulolytic activities showed significant positive correlation with microbial population, SMBC & SMBN.

**Table 5:** Impact of CA on available nutrient and microbial biomass based on ANOVA.

| Source of variance (SV) | df | Mean square |  |  |  |  |
| --- | --- | --- | --- | --- | --- | --- |
|  |  | SOC | Av. N | Av. P | SMBC | SMBN |
| Tillage (T) | 2 | 1.3383* | 236.14** | 24.36ns | 865.55* | 52.707ns |
| Error a | 2 | 0.0042 | 0.12 | 2.48 | 33.71 | 14.485 |
| Cropping system (CS) | 2 | 11.9110** | 615.30** | 371.48** | 775.64** | 268.289** |
| T × CS | 4 | 4.9568** | 350.38** | 20.50** | 396.17** | 20.024 ns |
| Error b | 6 | 0.0228 | 5.37 | 1.60 | 7.57 | 7.793 |
| RM (Residue Management) | 4 | 2.7683** | 197.39** | 23.60** | 914.43 ** | 33.316 ns |
| RM × T | 8 | 0.5165** | 31.80** | 14.02** | 494.28** | 23.762 ns |
| RM × CS | 8 | 0.9915** | 1.61** | 17.28** | 416.98** | 32.854 ** |
| RM × T × CS | 16 | 0.8852** | 27.08** | 9.09** | 198.27** | 37.305 ** |
| Error c | 36 | 0.0032 | 0.02 | 0.51 | 10.97 | 9.145 |
| CV (a) % |  | 0.6 % | 0.1 | 4.9 | 2.4 | 8.1 |
| CV (b) % |  | 1.5 % | 0.9 | 4.0 | 1.1 | 5.9 |
| CV (c) % |  | 0.6 % | 0.1 | 2.3 | 1.4 | 6.4 |
| Mean |  | 10.13 | 253.72 | 31.81 | 245.02 | 46.93 |

(ns: non-significant; ‘\*\*\*’ Significant at the 0.01 probability level; ‘\*’ Significant at the 0.05 probability level; df: degrees of freedom; CV: coefficient of variation)

**Table 6:**
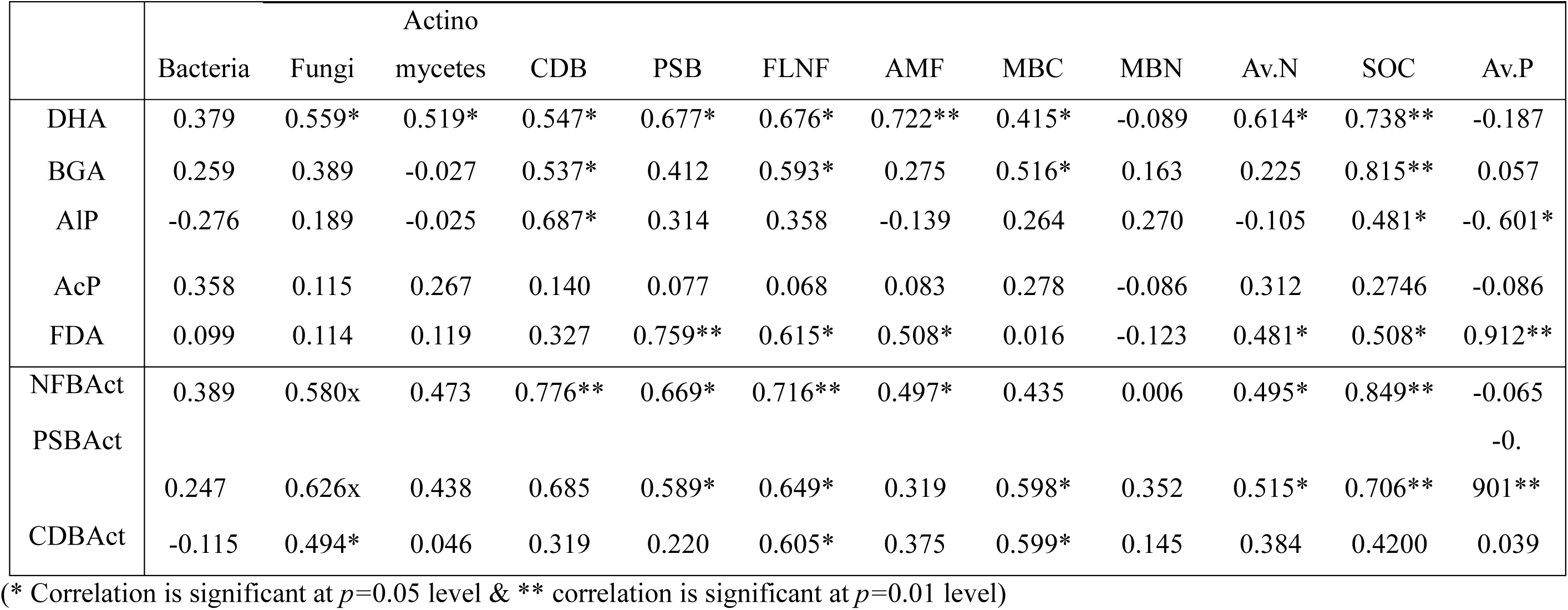
Correlation of soil enzyme and microbial activities with microbial and chemical properties of soil.

### 3.7 Ranking of different regimes of CA based Multivariate Analysis (MVA)

Ranking of practices under CA based on individual biochemical and microbial properties of soil showed following order of their suitability: ZT >RT >CT (Tillage), R_100_ > R_50_> R_0_ (degree of residue management) and RMCp ≈ RWGg >RCfBr (for CS). The outcome of ranking process indicates, for restoring microbial and biochemical properties of soil, CA is best management practices. Superior/elite conservation practices were identified by ranking through MVA, based on response of different biochemical and microbial properties of soil. The result indicated that irrespective of the CS, ZT or MT excelled when accompanied with R_100_. Higher ranks 1, 2, & 3 were attained by ZT-R_100_ under RWGg, MT-R_100_ under RWGg & ZT-R_100_ under RMCp, respectively. Contrarily to this, rank 25, 26 & 27 were associated with RCfBr under CT-R_50_, MT-R_0_ & CT-R_0_ (Table 7a). ZT, R_100_ & RWGg were best performer under the 27 combinations of CA. Total score under MVA indicated impact of treatments on biochemical and microbial properties of soil. Treatment combination with negative score were detrimental to soil biochemical and microbial properties while those with positive values indicated favorable treatment combination for soil properties studied. Treatment combinations CT-R_0_, MT-R_0_ & ZT-R_0_ under RMCp, CT-R_0_, CT-R_50_ & MT-R_0_ under RWGg and CT-R_0_, CT-R_50_, CT-R_100_, MT-R_0_, MT-R_50_ & ZT-R_0_ under RCfBr indicated negative score (Fig. 8a) while CT-R_0_, CT-R_50_, MT-R_0_, MT-R_50_, ZT-R_0_ & ZT-R_50_ under RMCp, CT-R_100_, MT-R_50_, MT-R_100_ and ZT-R_0_, ZT-R_50_ &ZT-R_100_ indicated positive score. Under RCfBr only treatments MT-R_100_, ZT-R_50_ & ZT-R_100_ showed positive value of total score.

**Table 7a:** Performance of conservation practices by ranking through multivariate analysis of biochemical and microbial properties of soil.

| CS | Tillage | Residue | PC1 | PC2 | Total Score | Rank |
| --- | --- | --- | --- | --- | --- | --- |
| RMCp | CT | R0 | -0.693 | 0.331 | -0.362 | 18 |
|  |  | R50 | 0.288 | -0.027 | 0.261 | 14 |
|  |  | R100 | 0.584 | 0.660 | 1.244 | 6 |
|  | MT | R0 | 0.162 | -1.053 | -0.890 | 20 |
|  |  | R50 | 0.581 | -0.182 | 0.399 | 12 |
|  |  | R100 | 0.758 | 0.194 | 0.951 | 8 |
|  | **ZT** | R0 | 0.069 | -1.321 | -1.252 | 21 |
|  |  | R50 | 0.871 | 0.645 | 1.517 | 4 |
|  |  | **R100** | 0.557 | 1.203 | **1.760** | **3** |
| RWGg | CT | R0 | 0.154 | -1.929 | -1.775 | 24 |
|  |  | R50 | 0.743 | -0.793 | -0.049 | 15 |
|  |  | R100 | 0.907 | -0.384 | 0.523 | 11 |
|  | MT | R0 | -0.177 | -1.459 | -1.636 | 23 |
|  |  | R50 | 0.855 | -0.076 | 0.779 | 9 |
|  |  | **R100** | 1.425 | 0.682 | 2.106 | **2** |
|  | **ZT** | R0 | 1.114 | -0.023 | 1.092 | 7 |
|  |  | R50 | 1.217 | -0.539 | 0.678 | 10 |
|  |  | **R100** | 1.420 | 1.205 | **2.625** | **1** |
| RCfBr | CT | R0 | -1.273 | -1.403 | -2.676 | 27 |
|  |  | R50 | -1.473 | -0.390 | -1.863 | 25 |
|  |  | R100 | -1.477 | 1.140 | -0.337 | 17 |
|  | MT | R0 | -1.488 | -1.001 | -2.489 | 26 |
|  |  | R50 | -1.034 | 0.319 | -0.715 | 19 |
|  |  | R100 | -0.678 | 0.984 | 0.306 | 13 |
|  | ZT | R0 | -1.705 | 0.213 | -1.492 | 22 |
|  |  | R50 | -0.663 | 0.557 | -0.106 | 16 |
|  |  | R100 | -1.045 | 2.446 | 1.401 | 5 |

**Fig 8a:**
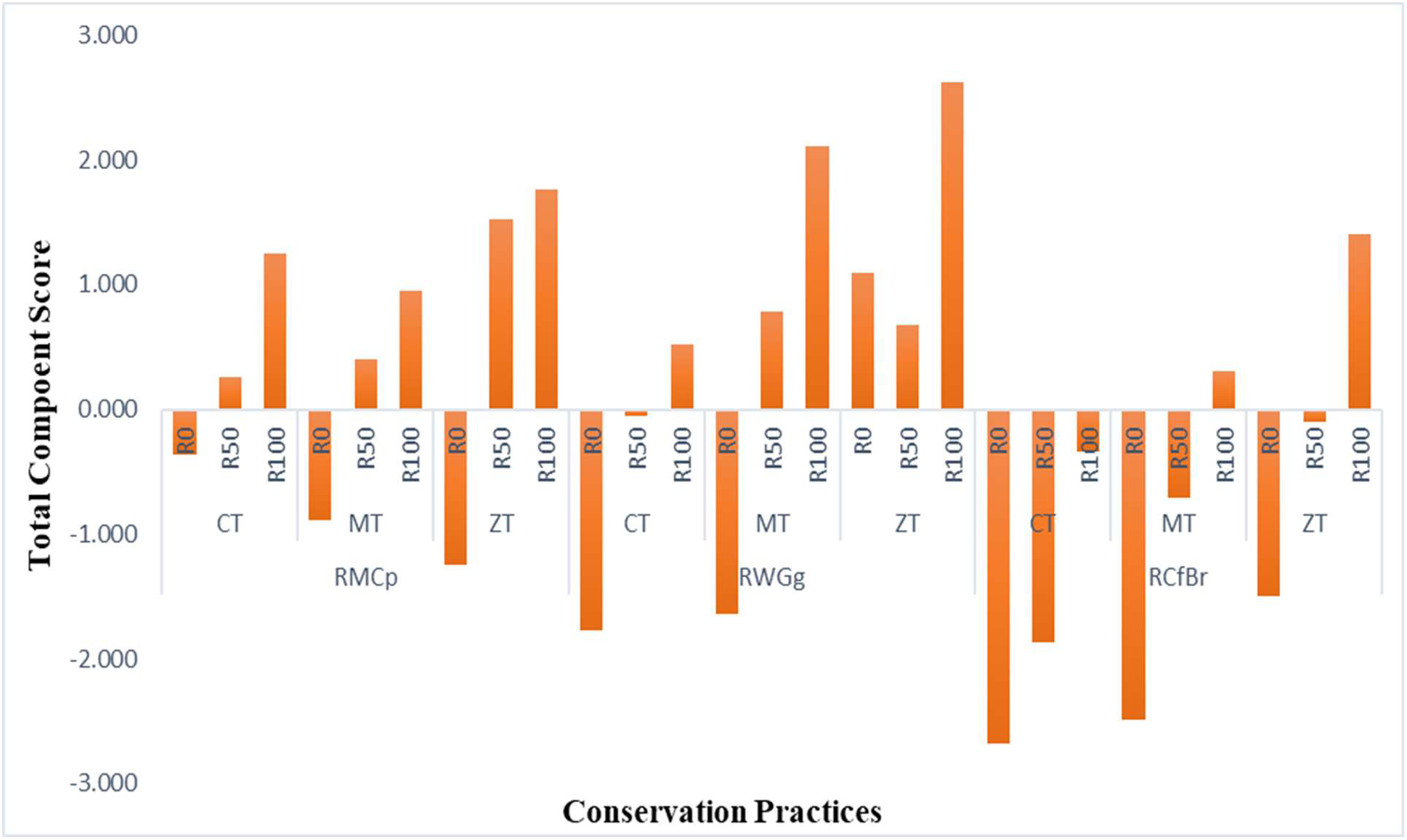
Total component score achieved by different conservation practices (where CT= Conventional tillage, MT= Minimum tillage and ZT= Zero tillage, R_0_: 0% Residue and 100% RDF, R_50_: 50% Residue and 100% RDF & 50% Residue and 75% RDF and R_100_:100% Residue and 50% RDF & 100% Residue and 75% RDF, RMCp: Rice-maize-cowpea, RWGg: Rice-wheat-green gram, RCfBr: Rice-cauliflower-bororice/summer rice)

### 3.8 Selection of minimum data set (MDS) of Biological Soil Health Index (BSHI)

The Statistics-based model was used to estimate soil quality index (SQI) using PCA. The PCA-model is used to create a minimum data set (MDS) to reduce the indicator load in the model and avoid data redundancy. All the observation (untransformed) physical, chemical and biological parameters (total 20) considered for SQI estimation was included in PCA. PC with eigen value (EV) >1 is chosen initially for screening of MDS. Additionally, first 7 PC considered, explained >80% variability in the dataset. The ‘highly weighted variables’ i.e., variables under a certain PC and absolute factor loading value within 10%, having highly weighted factor loading in component matrix is retained for MDS. The bold faces in component matrix factor loading (Table 7b) are retained for screening MDS. Finally, DHA, ACP, NFBAct, SOC, CDBAct, SMBN and SMBC were screened as MDS. After selection of parameters for the MDS, all selected observations were transformed using linear scoring functions (Fig 8b). The weightage for each parameter was calculated by the ratio between amount of variation explained by a particular PC and the maximum total variation in all PCs. BSHI was calculated using the equation:

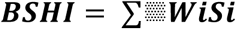

**Table 7b:** Factor loadings of different biochemical and microbial properties through multivariate PCA analysis.

| Parameters | PC1 | PC2 | PC3 | PC4 | PC5 | PC6 | PC7 |
| --- | --- | --- | --- | --- | --- | --- | --- |
| DHA | **0.882402** | -0.23115 | 0.047743 | -0.16503 | 0.026607 | 0.072895 | -0.00432 |
| BGA | 0.753556 | 0.067462 | 0.135916 | 0.295291 | -0.07797 | 0.266779 | -0.07434 |
| AlP | 0.415683 | 0.531969 | 0.460049 | 0.322528 | 0.075592 | -0.2471 | 0.110708 |
| Acp | 0.139636 | **-0.87941** | 0.198167 | 0.061285 | 0.119984 | -0.18053 | -0.02102 |
| FDA | 0.87937 | 0.065999 | -0.14745 | -0.2801 | 0.17126 | -0.00568 | -0.02803 |
| NFBAct | -0.14393 | 0.246453 | **0.615723** | -0.2941 | 0.487382 | -0.04813 | -0.06422 |
| PSBAct | 0.835942 | -0.12214 | -0.23954 | -0.10916 | 0.211447 | -0.11182 | -0.06713 |
| CDBAct | 0.540206 | -0.27289 | -0.06842 | 0.424349 | **0.497492** | -0.0022 | 0.115703 |
| Bacteria | 0.57583 | 0.632463 | 0.03636 | -0.01896 | 0.084488 | -0.13804 | 0.019287 |
| Fungi | 0.00108 | 0.860842 | 0.115751 | -0.10615 | 0.005514 | -0.16205 | 0.080214 |
| Actinomycetes | -0.56664 | 0.195558 | -0.15537 | -0.463 | 0.361091 | 0.01416 | 0.307775 |
| CDB | 0.508545 | 0.423062 | -0.45593 | 0.321478 | 0.114131 | 0.033301 | -0.33268 |
| PSB | -0.69141 | 0.432964 | -0.05071 | 0.142729 | 0.202918 | -0.16828 | -0.08031 |
| NFB | -0.50016 | 0.551738 | -0.52557 | -0.04065 | -0.01244 | -0.06509 | -0.21618 |
| AMF | 0.081684 | 0.841482 | -0.31015 | -0.02635 | 0.080871 | 0.21516 | 0.080863 |
| SOC | 0.431275 | -0.25651 | -0.01341 | **-0.54459** | -0.31226 | -0.23524 | -0.31958 |
| Available_N | 0.753651 | 0.394831 | -0.04137 | -0.29613 | 0.01391 | -0.19202 | 0.050476 |
| Available_P | 0.129398 | 0.648249 | 0.356681 | 0.189322 | -0.4508 | -0.25065 | 0.059148 |
| SMBC | 0.287215 | -0.29284 | -0.47617 | 0.080513 | -0.17948 | -0.32498 | **0.59986** |
| SMBN | -0.40131 | -0.32931 | -0.17384 | 0.192906 | 0.197878 | **-0.57687** | -0.2685 |

**Fig 8b:**
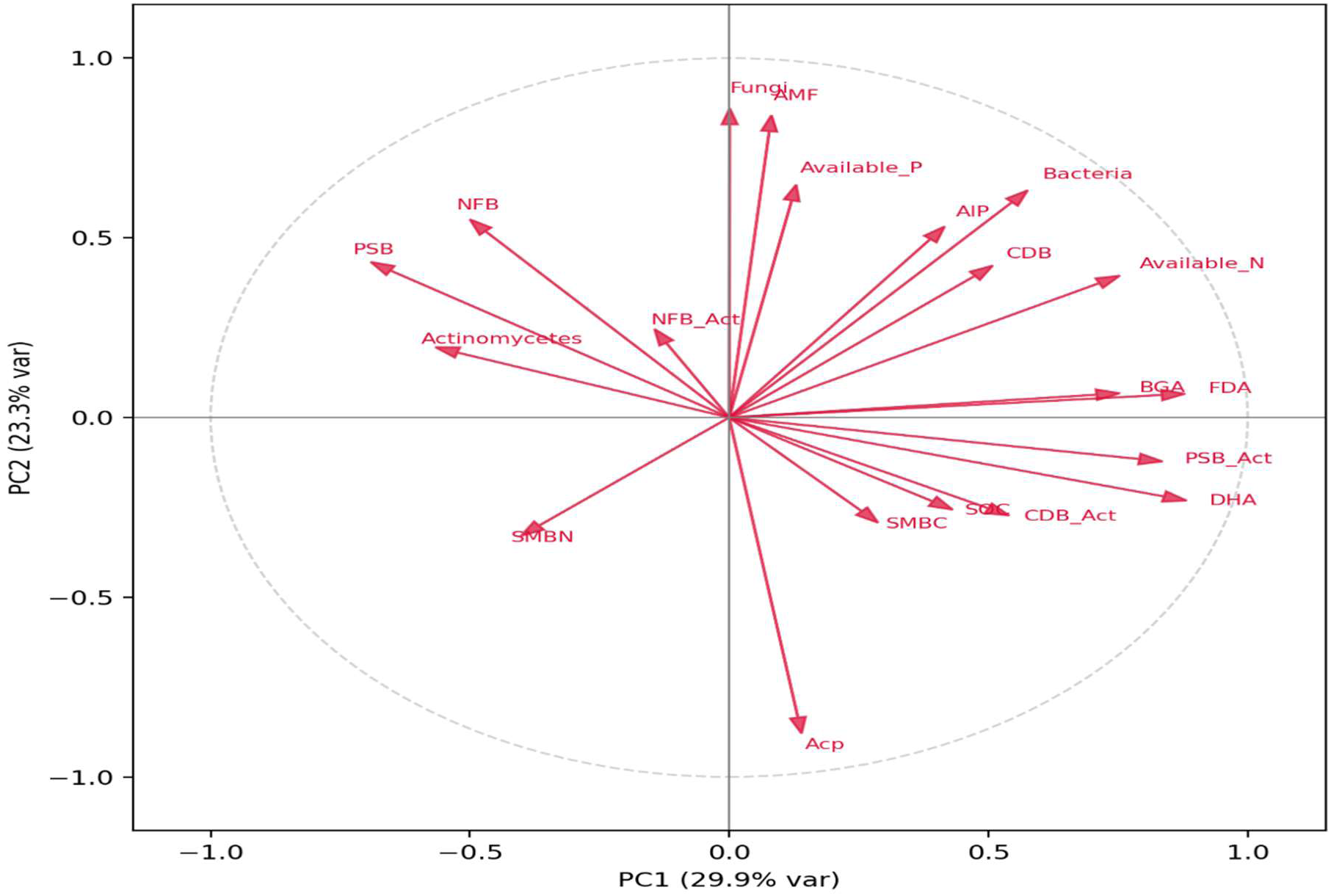
Vector biplot of parameters used for minimum dataset selection of BSHI.

Where, Wi = weight of MDS parameters, Si=linear scores of MDS parameters

Weighted SQI is computed as: ***BSHI= O.36 x DHA + 0.28 x AcP + 0.10 x NFB-Act + 0.08 x SOC + 0.07 x CDB-Act + 0.05 x SMBN + 0.05 x SMBC***

### 3.9 Effect of different treatments on BSHI

Analysis of the BSHI showed that, highest BSHI was observed in ZT followed by MT and CT but all the tillage practices were statistically at par with each other. Similarly, residue management levels (R_0_, R_50_, R_100_) were also at par but higher BSHI found in R_50_ & R_100_ over R_0_. In contrast, RCfBr achieved significantly higher BSHI values than RMCp and RWGg. Year effects were also distinct, with 2020 exhibiting a significantly higher BSHI than 2019 (Table 7c & Fig 8c).

**Table 7c:**
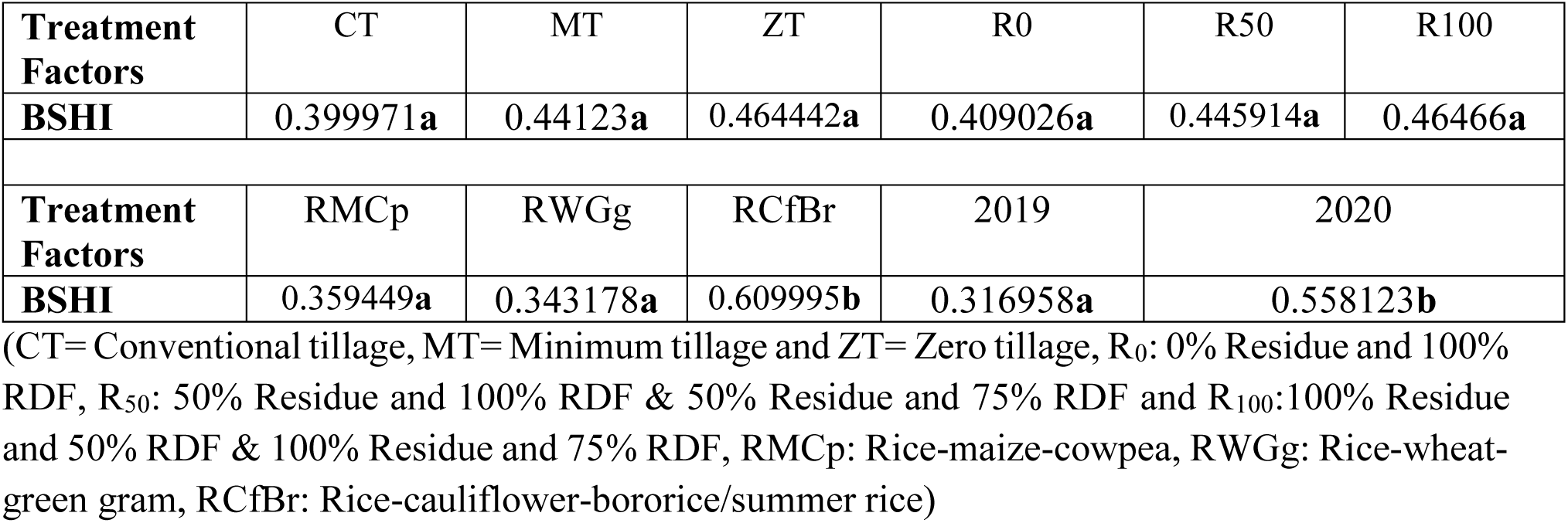
Biological soil health index for different treatments under CA.

| Treatment Factors | CT | MT | ZT | R<sub>0</sub> | R<sub>50</sub> | R<sub>100</sub> |
| --- | --- | --- | --- | --- | --- | --- |
| BSHI | 0.399971a | 0.44123a | 0.464442a | 0.409026a | 0.445914a | 0.46466a |
| Treatment Factors | RMCp | RWGg | RCfBr | 2019 | 2020 |  |
| BSHI | 0.359449a | 0.343178a | 0.609995b | 0.316958a | 0.558123b |  |

**Fig 8c:**
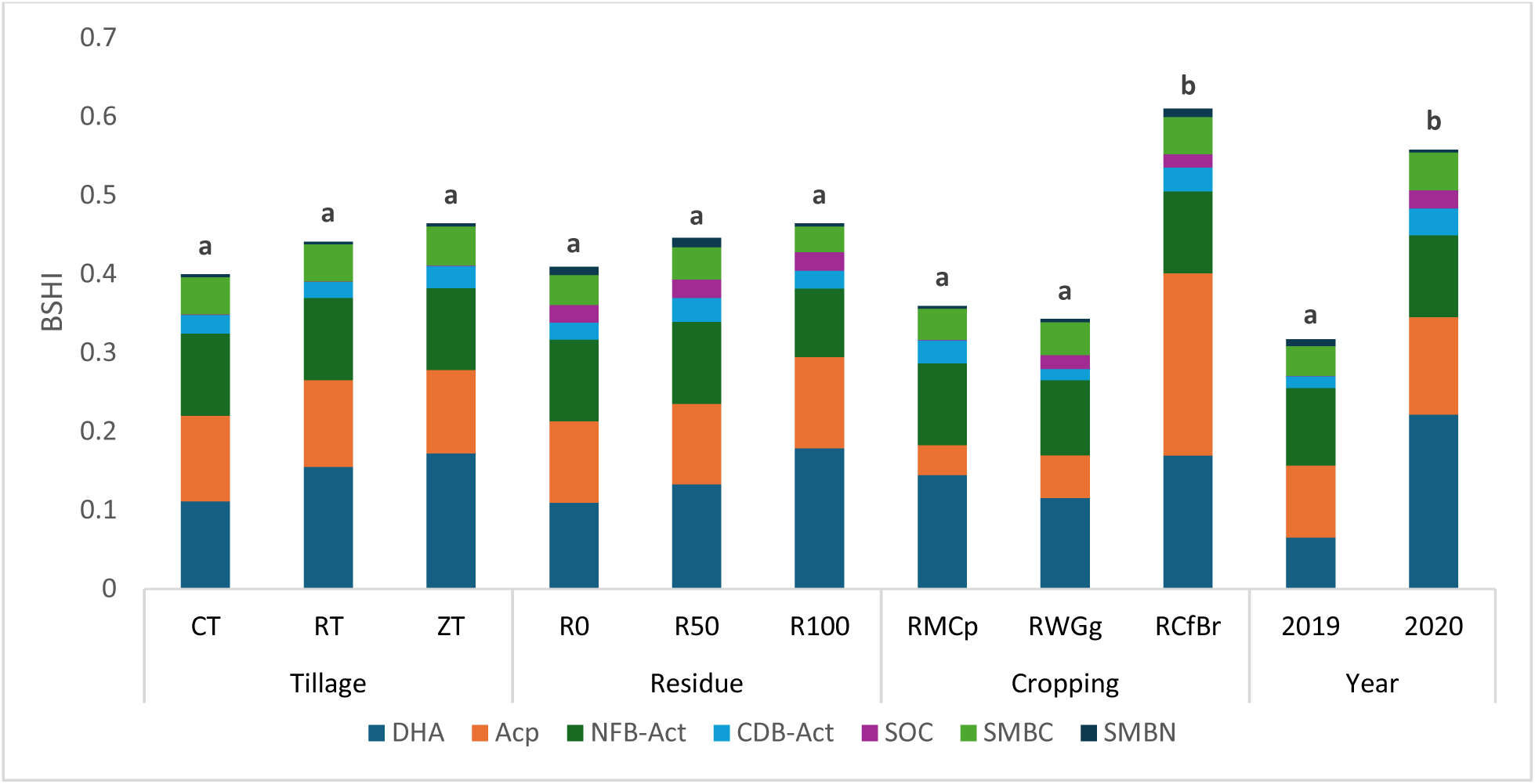
BSHI for different treatments under CA (where CT= Conventional tillage, MT= Minimum tillage and ZT= Zero tillage, R_0_: 0% Residue and 100% RDF, R_50_: 50% Residue and 100% RDF & 50% Residue and 75% RDF and R_100_:100% Residue and 50% RDF & 100% Residue and 75% RDF, NFBAct: Nitrogen fixation & CDBAct: Cellulolytic activities,SOC soil organic carbon, SMBC: soil microbial biomass carbon and SMBN: soil microbial biomass nitrogen

## 4. Discussion

Ortega et al. (2023) suggested ZT ensure lesser soil disturbance that protects SOM and SMB from mineralization, hence allowing better stratification ratio as well as quantity and quality of SOC preserved under ZT than CT, resulting to higher enzymatic activities, while conventional farming expose SOC to oxygen thus allow faster decomposition. Additionally, CA (ZT with residue) encourages macro-aggregate formation, especially within the topsoil by increasing SOC that protect soil microbes and enhance enzyme activity (Bhattacharyya et al. 2019). Whereas, mixing of soil in CT improve microbial contact with moisture & enzymes locked in different packets within soil aggregates, thus increase enzyme activities (Han et al. 2024). Aforementioned reasons may be due to varied impact of tillage on AlP under different CS (Fig 1). BGA under different tillage (i.e., extent of soil disturbance) may be associated with biochemical environments created, resulting to change in the populations of aerobic and facultative anaerobic microorganisms (Fig. 1c). Oxygen tension under ZT, reduce loss of added residue and produces water soluble low molecular weight phenolic compounds detrimental to microbes (Naresh et al. 2017) causing lower BGA.

High residue (R_100_) causes greater accumulation of SOC on the surface, increasing soil microbes and enzyme activities, especially BGA (De la Horra et al., 2003). Zhang et al. (2012) reported that microbial response is unaffected of residues incorporation ≤2.5 t/ha, however significant response occur at residue application 2.5-5 t/ha. 2018 onwards R_50_ and R_100_ received >2.5 t/ha residue, thus caused higher enzyme activities (Fig. 2c). Lower enzyme activities with higher fertilizer dose are associated with enzymes sensitivity towards chemical fertilizer. Higher nutrient concentration on fertilization enriches the soil with product of enzymes thus reduce their efficiency (Liang et al. 2014). Effects of crop rotation on enzyme action is associated with difference in root exudates, organic components from released from root systems and residues of crops under rotation which attract different microbial group to form unique rhizomicrobiome (Balota et al. 2004). Pulse enrich soil with nitrogen resulting to higher microbial activities than CS excluding legumes in rotation.

Higher microbial activities e.g., nitrogen fixation, phosphate solubilization, cellulose decomposition etc. in low input-farming e.g., CA are associated with better SOM accumulation, aggregate stability and microbial distribution (Pires et al. 2020), which led to improved nutrient uptake and plant growth (Bhardwaj et al. 2021). Reducing soil disturbance under MT/ZT can improve soil properties e.g., macropore structure, aggregate stability, nutrient availability which ultimately create suitable microbial habitat causing increase in microbial activities (Heidari et al. 2016). Whereas ploughing accelerate decomposition of SOM, modifies aggregate stability, and degrade soil quality (Lupwayi et al. 2012; Álvaro-Fuentes et al. 2013). Higher residue addition rejuvenate rhizosphere with diverse soil microbes, and carbon-rich substrates (Chinnadurai et al. 2014), resulting to higher nitrogen fixation, phosphorus solubilization and cellulolytic activities with higher residue application i.e., R_100_ ((Fig. 4 (i), (ii) & (iii)).

In all the three-CS where bacteria response was more prominent under residue application, fungi and actinomycetes population also showed remarkable response. As actinobacteria have more methyl-branched fatty acids; and methyl linoleate that typically belong to fungi, it reflects actinomycetes flourishment under harsh environment, Phospholipid fatty acid abundance was reported as nmol PLFA/g-soil. Soil microbes (bacteria, fungi, actinomycetes) are highly favoured in soil under maize & wheat, especially with residue applied as mulch under ZT. Residue incorporation add SOC making higher availability of food (i.e., C) to microorganisms, which, in turn, favours microbial growth and their biomass built up in soil. Fungi are more effectuated under ZT with residue mulching than bacteria. Fungi by virtue of their higher carbon utilization efficiency, flourish in undisturbed habitat and result to higher SMB (SMBC & SMBN) accumulation in soil (Singh et al. 2024). Residue protects soil, regulate its moisture & temperature, and prevent microbial exposure to harsh environment (De Quadros et al. 2012); enrich soil with C and N; maintains food availability (carbonaceous substrate) and provide energy for microbial growth, reproduction and produce higher biomass (Mangalassery et al. 2015), especially under cereals e.g., maize, and wheat.

SOC can be physically stabilized through the protection provided by soil aggregates under conservation tillage. Conservation tillage can substantially alter the size and distribution of soil aggregates (Gao et al. 2019a), and this may affect the accumulation of SOC. Organically fertilized soil shows higher OC hydrolases which decompose the added residues and along with constituting a good part of labile SOC, contributing to the higher oxidizable SOC (Xu et al. 2021). Different plant species used in the crop rotation (as cover crops or cash crop) release specific compounds in the soil and may cause change in the distribution of microbial communities (Chavarría et al. 2016). Pulse crops as endowed with inherent characteristics like BNF, release of root exudates and rhizodeposition of photosynthates, they are likely to change microbial properties of soil (Yusuf et al. 2009). Moreover, the biochemical compositions of legumes, low C:N ratio, higher ligno-polyphenols & protein content, rapid mineralization of residue and binding to the soil clay-complex (Sharma et al. 2021) would be the possible explanation of higher SOC in pulse-inclusive rotations. (Fig. 6).

Limited information is available regarding the impact of CA (residue and nutrient management under different CS) on N availability in soil (Mukherjee et al. 2024). Higher N availability under CA may be due to accumulation of carbonaceous materials in soils that enrich soil with potentially mineralizable N (PMN) fractions, which contribute nearly 40–60% organic N. PMN fractions are primarily composed of easily mineralizable protein or protein-like N compounds, these are vulnerable to quick mineralization for producing available N for plant uptake. Moreover, lignin, tannin, quinone bound protein-like compounds, amino sugars and other organic N complexing with polyvalent cations like Fe and Al may contribute to plant N availability in soil under CA (Mukherjee et al. 2024). As much as 90–95% of soil N is in organic forms and their dynamics and transformations largely depend on addition of organics residue under farming practices (Liu et al. 2018), hence available N was significantly higher in treatements with residue application than no residue application (Fig. 7). MT and residue are hypothesized to increase soil aggregation, and as a result, induce labile & organic P accumulation (Po), fasten its mineralization & reduce P sorption potential or fixation by Al, Fe and Ca by increasing anion competition for P binding sites (Johan et al. 2021). Better soil structure and aggregation in CA decreases the relative soil surface area for fixation and therefore potential sorption sites to which soluble P is exposed (Wei et al. 2017). Slower diffusion and P equilibration with sorption sites within larger aggregates reduce P fixation in weathered soils and ensure greater P availability (Margenot et al. 2017).

SMB is active fraction of SOC and highly sensitive to the changes in soil ecosystems due to management practices thus serve as an indicator of early changes in soil properties (Singh et al. 2024). Tillage practices and residue retention influence soil microclimate, distribution and decomposition of crop residue, and the mineralization and immobilization of nutrients (Cheng et al. 2017); such changes can alter SMB and microbial community structure (Li et al. 2018). In addition, cropping intensity alters the quality and quantity of C input into the soil thus it directly impacts SOC turnover and changes OC that could affect soil microbial community composition (Liu et al. 2018). SMBC was significantly higher under cereal-based cropping system with legumes in rotation compared to those excluding legumes while SMBN was higher in RWGg by 1.06 and 1.1 times than RMCp and RCfBr (Govaerts et al. 2007).

DHA is widely used to measure catabolic activities of soil microbes. DHA and AlP are associated with higher microbial activities and release of organic substances thereby creating “positive rhizosphere effect” including SMB (Chandra, 2011). Higher population of soil microbes is reflection of their activities in soil and positively correlated with DHA. SOC act as substrates for microbial proliferation and increased SMB, thereby increase enzymes activities (Borase et al. 2020). According to Liu et al. (2023) SOC is precursor of enzyme synthesis and play a vital role in their physical stabilization. Significant correlation of SOC & DHA, may be associated with microbial population and activities (Jha et al. 1992). Enzymes produced by soil microbes (FLNF, CDB etc) and organic substrate are active in top soil (Amer et al. 2017). Correlation of AlP with P is associated with its increased concentration in presence of phosphatase enzyme. Large amount of soil P exist in organic form (associated with esters & anhydrides of phosphoric acid), whereas plant prefer to take PO_4_^3-^. When P is limited in soil, plant roots and microbes increase the secretion of phosphatase to intensify P solubilization. Therefore, influencing the ability of the plant to cope with P-stressed conditions (Bautista and Ortiz, 2015). FDA is a potential soil quality indicator and show strong correlation with SMB, microbial populations (AMF, PSB, FLNF etc), DHA, AcP & AlP activity, SOC, and N and P content of the soil (Aseri & Tarafdar, 2006).

Cellulolytic microorganisms (genus *Streptomyces)* require nutrients to decompose added residue. If the soil is N deficient, they develop strategies to overcome low-N and sustain cellulolytic activities (Wang et al. 2021). *Streptomyces* are highly abundant cellulolytic microbes and secrete large number of hydrolytic enzymes that breakdown insoluble organic polymers, including chitin and cellulose into sugars/carbohydrates constituents. In addition, they act as highly efficient FLNF in soils and trigger the recycling of nutrients to maintain their cellulolytic process (Dahal et al. 2017). This multifunctionality of CDB might be the reason of significant correlation between CDB and BNF. Significant correlation of BNF with PSB and available P is associated with higher ATP formation in FLNF present in rhizosphere or bulk soil as conversion of N_2_ to NH_4_^+^ is highly energy intensive process as each molecule N_2_ fixation require 16 moles adenosine triphosphate (ATP) (Singh et al. 2024). SOM/SOC act as food to soil heterotrophs or mixotrophs and facilitate their growth, reproduction and N-fixing capacity of FLNF (Cheng et al. 2017). However, higher N (NH_4_^+^ and NO_3_^-^) in soil is inhibitory to FLNF as it represses the transcription and expression of nif-gene. In addition, many heterotrophic N_2_ fixers are facultative fixers and able to down-regulate their fixation pathway when other source of N is available in their ambient (Meng et al. 2012). In the N rich environment, microbes start uptake of inorganic N (NH_4_^+^ and NO_3_^-^), as it is less costly product in term of energy expenses than N fixation.

PSB activity ensure release of PO_4_^3-^ in soil from P containing compound. PO_4_^3-^ present in soil is used by microbes for synthesis of adenosine diphosphate (ADP) and ATP to perform different metabolic activities. This may cause flourishment of different soil microbes (e.g., fungi, FLNF, PSB, AMF etc.) (Billah et al. 2019). But significant negative correlation of soil P with PSB and PSBAct are associated with expression of *phoC* and *phoD* genes, part of the phosphate regulon (Pho) coding for AcP and AlP production, separately by PSB. Higher P has been observed to modulate (decrease) the expression of the *phoC* and *phoD* gene in soils. Organic fertilization e.g., residue application has been demonstrated to increase AlP and AcP activities in soil. However, *phoC*- and *phoD* harboring bacterial communities and their functional responses to chemical fertilization in acidic agricultural soils are not fully understood (Zheng et al. 2021).

Among the indicators selected for BSHI, DHA had the highest contribution. DHA being the key indicator of soil microbial activity and soil health as reflects its importance as the major parameter in providing the overview of metabolic state of the microbes. Acid phosphatases (AcP) and NFBAct were also found as important parameters for determining the BSHI, most probably due to their influence on N and C cycle within the soil environment which has overall impact on soil fertility and microbial growth. The rest of the parameter SOC, CBD-activity, SMBN and SMBC which are closely related to carbon metabolism by the soil microbes it justifies due to the presence of lot of residues and its usefulness as a representative of SOM decomposition. Higher degree of CA viz ZT-R_100_ tend to show higher BSHI values over conventional treatments but failed to be statistically significant this may be due to initial impact of CA on the soil systems. At the initial stage of CA adoption more recalcitrant C is retained in soil as root and straw residue which may have an immobilization effect, indicated by significantly lower BSHI in 2019 over 2020. The highest BSHI was found in 2020 which implies the systems tend to mineralize the excess C, it may also be the reason why the parameter in BSHI had many parameter which are indicators of soil C, N and P mineralization.

## 5. Conclusion

CA in lower IGP of West Bengal offer an appropriate agriculture practice for improving microbial and biochemical properties of soil. Such soil properties under conservation and conventional farming are captured distinctly. Study demonstrated that lesser soil disturbance, higher dose of residue addition and crop rotation with legumes, support microbial restoration, their biomass & activities and increase enzyme activities. Moreover, higher fertilization doses retard microbial activities by making availability of products. Residue addition should be recommended to improve soil microbial activity and thereby sustain the microbial environment of paddy soils and are key variables for maintaining the soil microbial and biochemical properties of the soil. The results are supported by statistical analysis like correlation, ANOVA, ranking through MVA etc. Treatments with higher intensity CA showed higher BSHI over convention treatments. Supporting data indicate improvement in soil characteristics, at initial stage (2-3 years) of CA adoption, hence more research work is needed to distinctly determine the possible result of long-term CA adoption on microbial and biochemical properties of soil.

## Acknowledgment

This research was carried out as a part of Ph.D. dissertation by the first author. The authors are thankful to the Indian Council of Agricultural Research, New Delhi for funding the work through the Centre for Advanced Agricultural Science and Technology on Conservation Agriculture under the National Agricultural Higher Education Project (Sanction No. NAHEP/CAAST/ 2017-18).

## References

A.T. Adetunji, F.B. Lewu, R. Mulidzi, B. Ncube, The biological activities of β-glucosidase, phosphatase and urease as soil quality indicators: a review. J. Soil Sci. Plant Nutr. 17 (2017) 794–807. 10.4067/S0718-95162017000300018

Á. Fuentes, F.J. Morell, E. Madejón, J. Lampurlanés, J.L. Arrúe, C. C. Martínez, Soil biochemical properties in a semiarid Mediterranean agroecosystem as affected by long-term tillage & N fertilization, Soil Till. Res. 129 (2013) 69–74. 10.1016/j.still.2013.01.005

A. Amer, B. Kashfa, and A. Bibi, Microbial β-glucosidase: sources, production and applications, J. Appl. & Environ. Microbiol. 5 (2017) 31–46. http://pubs.sciepub.com/jaem/5/1/4

S.S. Andrews, S. Susan, D. L. Karlen, and J. P. Mitchell, A comparison of soil quality indexing methods for vegetable production systems in Northern California, Agric. Ecosyst. Environ. 90 (2002) 25–45. 10.1016/S0167-8809(01)00174-8

G.K. Aseri, and J. C. Tarafdar, Fluorescein diacetate: a potential biological indicator for arid soils, Arid Land Res Manag. 20 (2006) 87–99. 10.1080/15324980500544473

P. Baldrian, and V. Valášková, Degradation of cellulose by basidiomycetous fungi, FEMS Microbiol. Rev. 32 (2008) 501–521. 10.1111/j.1574-6976.2008.00106.x

E. L. Balota, M. Kanashiro, A. C. Filho, D. S. Andrade, and R. P. Dick, Soil enzyme activities under long-term tillage and crop rotation systems in subtropical agro-ecosystems, Braz. J. Microbiol. 35 (2004) 300–306. 10.1590/S1517-83822004000300006

A. Cruz, Angélica, and Y. D. O. Hernández, Hydrolytic soil enzymes and their response to fertilization: a short review, Comun. Sci. 6 (2015) 255–262. 10.14295/cs.v6i3.962

A.K. Bhardwaj, D. Rajwar, R. K. Yadav, S.K. Chaudhari, D.K. Sharma, Nitrogen availability and use efficiency in wheat crop as influenced by the organic-input quality under major integrated nutrient management systems, Front. Plant Sci. 12 (2021) 634448. 10.3389/fpls.2021.634448

R. Bhattacharyya, T.K. Das, S. Das, A. Dey, A.K. Patra, R. Agnihotri, A. Ghosh, A.R. Sharma, Four years of conservation agriculture affects topsoil aggregate-associated 15nitrogen but not the 15nitrogen use efficiency by wheat in a semi-arid climate, Geoderma 337 (2019) 333–340. 10.1016/j.geoderma.2018.09.036

Billah, Motsim, M. Khan, A. Bano, T. U. Hassan, A. Munir, and A. R. Gurmani, “Phosphorus and phosphate solubilizing bacteria: Keys for sustainable agriculture.” Geomicrobiol. J. 36 (2019) 904–916. 10.1080/01490451.2019.1654043

Blake, G.R. & K. H. Hartge, Bulk density in A. Klute (Eds.) Methods of soil analysis: Part 1 Physical and mineralogical methods, Wiley Online Library, Soil Science Society of America, 1986, pp. 363–375. 10.2136/sssabookser5.1.2ed.c13

Borase, D. N., C. P. Nath, K. K. Hazra, M. Senthilkumar, S. S. Singh, C. S. Praharaj, U. Singh, and N. Kumar. Long-term impact of diversified crop rotations and nutrient management practices on soil microbial functions and soil enzymes activity, Ecol. Indic. 114 (2020) 106322. 10.1016/j.ecolind.2020.106322

J J. Brejda, T. B. Moorman, D. L. Karlen, T. H. Dao, “Identification of regional soil quality factors and indicators I. Central and Southern High Plains.” Soil Sci. Soc. Am. J. 64 (2000) 2115–2124. 10.2136/sssaj2000.6462115x

A. Ramesh, “Effect of summer crops and their residue management on yield of succeeding wheat and soil properties. J. Indian Soc. Soil Sci, 59 (2011) 37.

B. N. Chavarría, R.A. Verdenelli, D.L. Serri, S.B. Restovich, A.E. Andriulo, J.M. Meriles, S. V. Gil, Effect of cover crops on microbial community structure and related enzyme activities and macronutrient availability, Eur. J. Soil Biol. 76 (2016) 74–82. 10.1016/j.ejsobi.2016.07.002

Yi Cheng, J. Wang, J. Wang, S. X. Chang, S. Wang, The quality and quantity of exogenous organic carbon input control microbial NO3− immobilization: a meta-analysis.” Soil Biol. Biochem. 115 (2017) 357–363. 10.1016/j.soilbio.2017.09.006

C. Chinnadurai, G. Gopalaswamy, and D. Balachandar, Long term effects of nutrient management regimes on abundance of bacterial genes and soil biochemical processes for fertility sustainability in a semi-arid tropical Alfisol, Geoderma 232 (2014) 563–572. 10.1016/j.geoderma.2014.06.015

B. Dahal, G. NandaKafle, L. Perkins, and V. S. Brözel, Diversity of free-Living nitrogen fixing Streptomyces in soils of the badlands of South Dakota, Microbiol. Res. 195 (2017) 31–39. 10.1016/j.micres.2016.11.004

De la Horra, A. M., M. E. Conti, and R. M. Palma, β-glucosidase and proteases activities as affected by long-term management practices in a Typic Argiudoll soil, Commun. Soil Sci. Plant Anal. 34.17-18 (2003) 2395–2404. 10.1081/CSS-120024775

P. D. de Quadros, K. Zhalnina, A. D. Richardson, J. R. Fagen, J. Drew, C. Bayer, F.A.O. Camargo, and E. W. Triplett, The effect of tillage system and crop rotation on soil microbial diversity and composition in a subtropical acrisol, Diversity. 4 (2012) 375–395. 10.3390/d4040375

R. P. Dick, D. P. Breakwell, and R. F. Turco, Soil enzyme activities and biodiversity measurements as integrative microbiological indicators in J. W. Doran, A. J. Jones (Eds.), Methods for assessing soil quality, Soil Science Society of America, 1997, pp. 247–271. 10.2136/sssaspecpub49.c15

R. P. Dick, P. E. Rasmussen & E. A. Kerle, Influence of long-term residue management on soil enzyme activities in relation to soil chemical properties of a wheat-fallow system. Biol. Fertil. Soils, 6 (1988) 159–164. 10.1007/BF00257667

F. Eivazi, and M. A. Tabatabai, Phosphatases in soils, Soil Biol. Biochem. 9(3) (1977) 167–172. 10.1016/0038-0717(77)90070-0

F. Eivazi, & M. A. Tabatabai, Glucosidases and galactosidases in soils, Soil Biol. Biochem. 20(5) (1988) 601–606. 10.1016/0038-0717(88)90141-1

L. Gao, B. Wang, S. Li, Y. Han, X. Zhang, D. Gong, M. Ma, G. Liang, H. Wu, X. Wu, D. Cai, A. Degré, Effects of different long-term tillage systems on the composition of organic matter by 13C CP/TOSS NMR in physical fractions in the Loess Plateau of China, Soil Till. Res. 194 (2019) 104321. 10.1016/j.still.2019.104321

B. Govaerts, M. Mezzalama, Y. Unno, K. D. Sayre, M. Luna-Guido, K. Vanherck, & J. Deckers, Influence of tillage, residue management, and crop rotation on soil microbial biomass and catabolic diversity. Appl. Soil Ecol, 37 (1-2) (2007) 18–30. 10.1016/j.apsoil.2007.03.006

J. Habig, & C. Swanepoel, Effects of conservation agriculture and fertilization on soil microbial diversity and activity. Environ. 2(3) (2015) 358–384. https://www.mdpi.com/2076-3298/2/3/358

C. Han, W. Zhou, Y. Gu, J. Wang, Y. Zhou, Y. Xue, & K. H. Siddique, Effects of tillage regime on soil aggregate-associated carbon, enzyme activity, and microbial community structure in a semiarid agroecosystem, Plant Soil. 498(1) (2024) 543–559. 10.1007/s11104-023-06453-1

J.J. Hanway, & H. Heidel, (1952). Soil analysis methods as used in Iowa State. J. Iowa Agri. 57 (1952), 57:1–131

G. Heidari, K. Mohammadi, & Y. Sohrabi, Responses of soil microbial biomass and enzyme activities to tillage and fertilization systems in soybean (Glycine max L.) production. Front. Plant Sci. 7 (2016), 1730. 10.3389/fpls.2016.01730

B. Hutter, & T. Dick, Increased alanine dehydrogenase activity during dormancy in Mycobacterium smegmatis. FEMS Microbiol. Lett. 167(1) (1998) 7–11. 10.1111/j.1574-6968.1998.tb13200.x

M. Kumar, S.K. Singh, P., Raina, B. K Sharma, Status of available major and micronutrients in arid soils of Churu district of western Rajasthan. J. Indian Soc. Soil Sci. 59(2), (2011) 188

D. S. Jenkinson, The effects of biocidal treatments on metabolism in soil—IV. The decomposition of fumigated organisms in soil. Soil Bio. Biochem. 8(3) (1976) 203–208. 10.1016/0038-0717(76)90004-3

D.K. Jha, G. D. Sharma, & R. R. Mishra, Soil microbial population numbers and enzyme activities in relation to altitude and forest degradation. Soil Bio. Biochem. 24(8) (1992) 761–767. 10.1016/0038-0717(92)90250-2

P. D. Johan, O. H. Ahmed, L. Omar, & N. A. Hasbullah, Phosphorus transformation in soils following co-application of charcoal and wood ash. Agronomy, 11(10) (2010) 10.3390/agronomy11102010

O. A. Leal, T. J. Amado, J. E. Fiorin, C. Keller, G. B. Reimche, C. W. Rice, & R. Schwalbert, Linking cover crop residue quality and tillage system to CO2-C emission, soil C and N stocks and crop yield based on a long-term experiment. Agronomy, 10(12) (2020) 1848. 10.1080/00207233.2018.1494927

Y. Li, S. X. Chang, L. Tian, & Q. Zhang, Conservation agriculture practices increase soil microbial biomass carbon and nitrogen in agricultural soils: A global meta-analysis. Soil Bio. Biochem. 121 (2018) 50–58. 10.1016/j.soilbio.2018.02.024

Q. Liang, H. Chen, Y. Gong, H. Yang, M. Fan, & Y. Kuzyakov, Effects of 15 years of manure and mineral fertilizers on enzyme activities in particle-size fractions in a North China Plain soil. Eur. J. Soil Biol. 60 (2014) 112–119. 10.1016/j.ejsobi.2013.11.009

Q. Liu, Y. Zhang, B. Liu, J. E. Amonette, Z. Lin, G. Liu, & Z. Xie, How does biochar influence soil N cycle? A meta-analysis. Plant Soil, 426 (2018) 211–225. 10.1007/s11104-018-3619-4

X. Liu, X. Song, S. Li, G. Liang, & X. Wu, Understanding how conservation tillage promotes soil carbon accumulation: Insights into extracellular enzyme activities and carbon flows between aggregate fractions. Sci. Total Environ. 897 (2023) 165408. 10.1016/j.scitotenv.2023.165408

N. Z. Lupwayi, G. P. Lafond, N. Ziadi, & C. A. Grant, Soil microbial response to nitrogen fertilizer and tillage in barley and corn. Soil Tillage Res. 118 (2012) 139–146. 10.1016/j.still.2011.11.006

S. Mangalassery, S. J. Mooney, D. L. Sparkes, W. T. Fraser, & S. Sjögersten, Impacts of zero tillage on soil enzyme activities, microbial characteristics and organic matter functional chemistry in temperate soils. Eur. J. Soil Biol. 68 (2015) 9–17. 10.1016/j.ejsobi.2015.03.001

A. J. Margenot, B. K. Paul, R. R. Sommer, M. M. Pulleman, S. J. Parikh, L. E. Jackson, & S. J. Fonte, Can conservation agriculture improve phosphorus (P) availability in weathered soils? Effects of tillage and residue management on soil P status after 9 years in a Kenyan Oxisol. Soil Tillage Res. 166 (2017) 157–166. 10.1016/j.still.2016.09.003

I. C. Mendes, D. M. G. Sousa, O. D. Dantas, A. A. C. Lopes, F. B. R. Junior, M. I. Oliveira, & G. M. Chaer, Soil quality and grain yield: A win–win combination in clayey tropical Oxisols. Geoderma, 388 (2021) 114880. 10.1016/j.geoderma.2020.114880

X. Meng, L. Wang, X. Long, Z. Liu, Z. Zhang, & Zed, R. Influence of nitrogen fertilization on diazotrophic communities in the rhizosphere of the Jerusalem artichoke (Helianthus tuberosus L.). Res. Microbio. 163(5) (2012) 349–356. 10.1016/j.resmic.2012.03.005

S. Mukherjee, D. Sarkar, B. Mandal, S. Kanthal, S. Ghosh, B. Sahu,….& N. Saha, Conservation agriculture influences soil nitrogen availability in the lower Indo-Gangetic Plains. Plant Soil, 508 (2024) 1–17. 10.1007/s11104-024-06826-0

R. K. Naresh, A. S. Panwar, S. S. Dhaliwal, R. K. Gupta, A. Kumar, R. S. Rathore, & N. C. Mahajan, Effect of organic inputs on strength and stability of soil aggregates associated organic carbon concentration under rice-wheat rotation in Indo-Gangetic Plain zone of India. Int. J. Curr. Microbiol. Appl. Sci, 6(10) (2017) 1973–2008. 10.20546/ijcmas.2017.610.237

S. R. Olsen, Estimation of available phosphorus in soils by extraction with sodium bicarbonate (No. 939). US Department of Agriculture, 1954

R. Ortega, I. Miralles, R. Soria, N. Rodríguez-Berbel, A. B. Villafuerte, D. A. Zema, & M. E. Lucas-Borja, Short-term effects of post-fire soil mulching with wheat straw and wood chips on the enzymatic activities in a Mediterranean pine forest. Sci. Total Environ. 857 (2023) 159489. 10.1016/j.scitotenv.2022.159489

C. M., Parihar, M. R. Yadav, S. L. Jat, A. K. Singh, B. Kumar, S. Pradhan,… & O. P. Yadav, Long term effect of conservation agriculture in maize rotations on total organic carbon, physical and biological properties of a sandy loam soil in north-western Indo-Gangetic Plains. Soil Tillage Res. 161 (2016) 116–128. 10.1016/j.still.2016.04.001

J. H. Passinato, T. J. Amado, A. Kassam, J. A. Acosta, & L. D. P. Amaral, Soil health check-up of conservation agriculture farming systems in Brazil. Agronomy, 11(12) (2021) 2410. 10.3390/agronomy11122410

M. B. Peoples, & E. T. Craswell, Biological nitrogen fixation: investments, expectations and actual contributions to agriculture. Plant Soil, 141 (1992) 13–39. 10.1007/BF00011308

A. C. A. Pires, T. J. Amado, G. Reimche, R. Schwalbert, M. V. Sarto, R. S Nicoloso, & B. W. Rice, Diversified crop rotation with no-till changes microbial distribution with depth and enhances activity in a subtropical Oxisol. Eur. J. Soil Sci. 71(6) (2020) 1173–1187. 10.1111/ejss.12981

A. E. Richardson, Prospects for using soil microorganisms to improve the acquisition of phosphorus by plants. Funct. Plant Biol. 28(9) (2001) 897–906. 10.1071/PP01093

Saha, N., Mandal, B. Soil Health – A Precondition for Crop Production. In: Khan, M., Zaidi, A., Musarrat, J. (eds) Microbial Strategies for Crop Improvement. Springer, Berlin, Heidelberg. 2009 10.1007/978-3-642-01979-1_8

S. Sharma, R. Saikia, H. S. Thind, Y. Singh, & M. L. Jat, Tillage, green manure and residue management accelerate soil carbon pools and hydrolytic enzymatic activities for conservation agriculture-based rice-wheat systems. Commun Soil Sci Plant Anal. 52(5) (2021) 470–486. 10.1080/00103624.2020.1862147

P. Singh, S. Dutta, S. Mukherjee, N. Saha, B. Dash, S. Ghosh, …. & B. Mandal, Spatiotemporal Shift of Soil Microbes in Conservation Agriculture under a Rice-Based Cropping System at the New Alluvial Zone of Lower Gangetic Plain. J. Soil Sci. Plant Nutr. 24 (2024) 1–15. 10.1007/s42729-024-01785-y

P. Singh, S. Mukherjee, N. Saha, S. Biswas, & B. Mandal, Conservation Agriculture in Reshaping Belowground Microbial Diversity. In: Rakshit, A., Singh, S., Abhilash, P., Biswas, A. (eds) Soil Science: Fundamentals to Recent Advances. Springer, Singapore, 2021 https://link.springer.com/chapter/10.1007/978-981-16-0917-6_8

A. V. Subbiah, & G. L. Asija, A rapid procedure for the estimation of available nitrogen in soils. Curr Sci 25 (1956) 259–260

A. Walkley, and I.A. Black An examination of the Degtjareff method for determining soil organic matter and a proposed modification of the chromic acid titration method. Soil Sci. 37 (1934) 29–38

H. L. Wang, Z. K. Gong, J. H. Wu, D.L Wang, Cellulolytic bacteria capable of nitrogen fixation in saline-sodic grassland soils. Appl. ecol. environ. res. 19(2) (2021) 10.15666/aeer/1902_867879

K. Wei, H. Bao, S. Huang, & L. Chen, Effects of long-term fertilization on available P, P composition and phosphatase activities in soil from the Huang-Huai-Hai Plain of China. Agric. Ecosyst. Environ. 237 (2017) 134–142. 10.1016/j.agee.2016.12.030

A. Wittmann, K. Riedel, & R. D. Schmid, Microbial and enzyme sensors for environmental monitoring. In Handbook of biosensors and electronic noses CRC Press 2024. pp. 299–332.

H. Xu, Q. Qu, Y. Chen, G. Liu, & S. Xue, Responses of soil enzyme activity and soil organic carbon stability over time after cropland abandonment in different vegetation zones of the Loess Plateau of China. Catena, 196 (2021) 104812. 10.1016/j.catena.2020.104812

A. A. Yusuf, R. C. Abaidoo, E. N. O. Iwuafor, O. O. Olufajo, & N. Sanginga, Rotation effects of grain legumes and fallow on maize yield, microbial biomass and chemical properties of an Alfisol in the Nigerian savanna. Agric. Ecosyst. Environ, 129 (1-3) (2009) 325–331. 10.1016/j.agee.2008.10.007

Q. C. Zhang, I. H. Shamsi, D. T. Xu, G. H. Wang, X. Y. Lin, G. Jilani, & A. N. Chaudhry, Chemical fertilizer and organic manure inputs in soil exhibit a vice versa pattern of microbial community structure. Appl. Soil Ecol. 57 (2012) 1–8. 10.1016/j.apsoil.2012.02.012

M. M. Zheng, C. Wang, W. X. Li, L. Guo, Z. J. Cai, B. R. Wang, & R. F. Shen, Changes of acid and alkaline phosphatase activities in long-term chemical fertilization are driven by the similar soil properties and associated microbial community composition in acidic soil. Eur. J. Soil Biol. 104 (2021) 103312. 10.1016/j.ejsobi.2021.103312

S.M. Zuber, and M.B. Villamil, Meta-analysis approach to assess effect of tillage on microbial biomass and enzyme activities. Soil Bio. Biochem. 97 (2016) 176–187.

